# Discovery of an Antarctic ascidian-associated uncultivated *Verrucomicrobia* with antimelanoma palmerolide biosynthetic potential

**DOI:** 10.1101/2021.05.05.442870

**Authors:** Alison E. Murray, Chien-Chi Lo, Hajnalka E. Daligault, Nicole E. Avalon, Robert W. Read, Karen W. Davenport, Mary L. Higham, Yuliya Kunde, Armand E.K. Dichosa, Bill J. Baker, Patrick S.G. Chain

**Affiliations:** Division of Earth and Ecosystem Science, Desert Research Institute, Reno, Nevada, USA; Bioscience Division, Los Alamos National Laboratory, Los Alamos, New Mexico, USA; Department of Chemistry, University of South Florida, Tampa, Florida, USA

## Abstract

The Antarctic marine ecosystem harbors a wealth of biological and chemical innovation that has risen in concert over millennia since the isolation of the continent and formation of the Antarctic circumpolar current. Scientific inquiry into the novelty of marine natural products produced by Antarctic benthic invertebrates led to the discovery of a bioactive macrolide, palmerolide A, that has specific activity against melanoma and holds considerable promise as an anticancer therapeutic. While this compound was isolated from the Antarctic ascidian *Synoicum adareanum*, its biosynthesis has since been hypothesized to be microbially mediated, given structural similarities to microbially-produced hybrid non-ribosomal peptide-polyketide macrolides. Here, we describe a metagenome-enabled investigation aimed at identifying the biosynthetic gene cluster (BGC) and palmerolide A-producing organism. A 74 Kbp candidate BGC encoding the multi-modular enzymatic machinery (hybrid Type I-*trans*-AT polyketide synthase-non-ribosomal peptide synthetase and tailoring functional domains) was identified and found to harbor key features predicted as necessary for palmerolide A biosynthesis. Surveys of ascidian microbiome samples targeting the candidate BGC revealed a high correlation between palmerolide-gene targets and a single 16S rRNA gene variant (R=0.83 – 0.99). Through repeated rounds of metagenome sequencing followed by binning contigs into metagenome-assembled genomes, we were able to retrieve a near-complete genome (10 contigs) of the BGC-producing organism, a novel verrucomicrobium within the *Opitutaceae* family that we propose here as *Candidatus* Synoicihabitans palmerolidicus. The refined genome assembly harbors five highly similar BGC copies, along with structural and functional features that shed light on the host-associated nature of this unique bacterium.

**Importance:** Palmerolide A has potential as a chemotherapeutic agent to target melanoma. We interrogated the microbiome of the Antarctic ascidian, *Synoicum adareanum*, using a cultivation-independent high-throughput sequencing and bioinformatic strategy. The metagenome-encoded biosynthetic machinery predicted to produce palmerolide A was found to be associated with the genome of a member of the *S. adareanum* core microbiome. Phylogenomic analysis suggests the organism represents a new deeply-branching genus, *Candidatus* Synoicihabitans palmerolidicus, in the *Opitutaceae* family of the Verrucomicrobia phylum. The *Ca*. S. palmerolidicus 4.29 Mb genome encodes a repertoire of carbohydrate-utilizing and transport pathways enabling its ascidian-associated lifestyle. The palmerolide-producer’s genome also contains five distinct copies of the large palmerolide biosynthetic gene cluster that may provide structural complexity of palmerolide variants.

## Introduction

Across the world’s oceans, marine benthic invertebrates harbor a rich source of natural products that serve metabolic and ecological roles in situ. These compounds provide a multitude of medicinal and biotechnological applications to science, health and industry. The organisms responsible for their biosynthesis are often unknown (1, 2). Increasingly, these metabolites, especially in the polyketide class (*trans*-AT in particular), are found to be produced by microbial counterparts associated with the invertebrate host (3-5). Invertebrates including sponges, corals, and ascidians for example, harbor a wealth of diverse microbes, few of which have been cultivated (e.g., 6-8). Genomic tools, in particular, are revealing biochemical pathways potentially critical in the host-microbe associations (9). Microbes that form persistent mutualistic (symbiotic) associations provide key roles in host ecology, such as provision of metabolic requirements, production of adaptive features such as photoprotective pigments, bioluminescence, or antifoulants, and biosynthesis of chemical defense agents.

Antarctic marine ecosystems harbor species-rich macrobenthic communities (10-12), which have been the subject of natural products investigations over the past 30 years resulting in the identification of > 600 metabolites (13). Initially, it was not known whether the same selective pressures (namely predation and competition, e.g., 14) that operate in mid and low latitudes would drive benthic organisms at the poles to create novel chemistry (15). However, this does appear to be the case, and novel natural products have been discovered across algae, sponges, corals, nudibranchs, echinoderms, bryozoans, ascidians, and increasingly amongst microorganisms (16) for which the ecological roles have been deduced in a number of cases (13, 17). Studies of Antarctic benthic invertebrate-microbe associations however, pale in comparison to studies at lower latitudes, yet the few studies that have been reported suggest these associations (i) harbor an untapped reservoir of biological diversity (18-21) including fungi (22), (ii) are host species-specific (23, 24), (iii) provide the host with sources of nitrogen and fixed carbon (25) and (iv) have biosynthetic functional potential (26, 27).

This study was specifically motivated by our desire to understand the biosynthetic origins of a natural product, palmerolide A, given its potent anticancer activity (28), that is found to be associated with the polyclinid Antarctic ascidian, *Synoicum adareanum* (Fig. 1A and B). Ascidians are known to be rich sources of bioactive natural products (9). They have been found to harbor polyketide, terpenoid, peptide, alkaloid and a few other classes of natural products, of which the majority have cytotoxic and/or antimicrobial activities. In addition to palmerolide A, a few other natural products derived from Antarctic ascidians have been reported (29-31).

**Figure 1.**
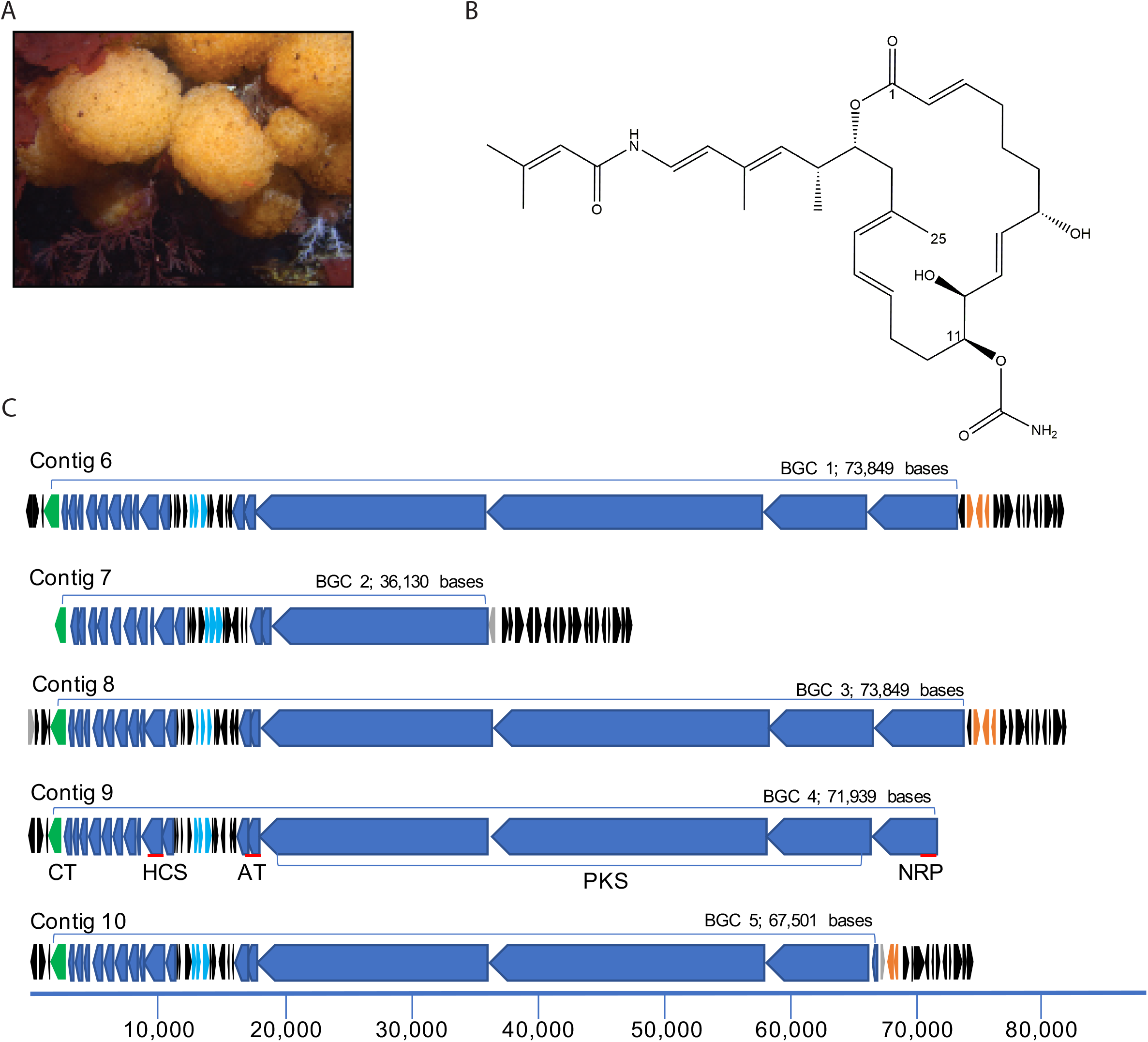
Palmerolide A, a cytotoxic, macrolide with anti-melanoma activity is found in the tissues of *Synoicum adareanum* in which a candidate biosynthetic gene cluster has been identified. (A) *S. adareanum* occurs on rocky coastal seafloor habitats in the Antarctic; this study focused on the region off-shore of Anvers Island in the Antarctic Peninsula. (B) Palmerolide A, is the product of a hybrid PKS-NRPS system in which biosynthesis begins with a PKS starter unit followed by incorporation of a glycine subunit by an NRPS module. Subsequent elongation, cyclization and termination steps follow. Two additional features of the molecule include a methyl group on C-25 and a carbamate group on C-11. (C) Five repeats encompassing candidate palmerolide biosynthetic gene clusters were identified. The BGC (in blue) is defined as starting with the NRP unit and ending at the carbamoyltransferase (green). Candidate palmerolide A biosynthetic gene cluster BGC4 was identified from initial metagenome library assemblies. The other four clusters were identified following a third round of sequencing, assembly and manual finishing. Primary BGC coding sequences (CDS) and a conserved tailoring cassette are in blue. Light blue CDS are an ATP transporter with homology to an antibiotic transporter, SyrD. All black CDS are repeated amongst the BGCs. Orange CDS are transposase/integrase domains. Gray CDS are unique, non-repeated; and in BGC2 and 5, the unique CDS encode transposases, distinct from the predicted amino acid sequences of those in orange. The red lines associated with Contig 9 indicate targeted quantitative PCR regions.

Ascidian-associated microbes responsible for natural product biosynthesis have been shown to be affiliated with bacterial phyla including Actinobacteria (which dominates the recognized diversity), Cyanobacteria, Firmicutes, Proteobacteria (both Alphaproteobacteria and Gammaproteobacteria) and Verrucomicrobia in addition to many fungi (32, 33). Metagenome-enabled studies have been key in linking natural products to the organisms producing them in a number of cases, e.g. patellamide A and C to Cyanobacteria-affiliated *Prochloron* spp. (4), the tetrahydroisoquinoline alkaloid ET-743 to Gammaproteobacteria-affiliated *Candidatus* Endoecteinascidia frumentensis (34), patellazoles to Alphaproteobacteria-affiliated *Candidatus* Endolissoclinum faulkneri (35), and mandelalides to Verrucomicrobia-affiliated *Candidatus* Didemnitutus mandela (36). However, this is most certainly an under-representation of the diversity of ascidian-associated microorganisms with capabilities for synthesizing bioactive compounds, given the breadth of ascidian biodiversity (37). These linkages have been yet to be investigated for Antarctic ascidians.

Palmerolide A has anticancer properties with selective activity against melanoma when tested in the National Cancer Institute 60 cell-line panel (28). This result is of particular interest, as there are few natural product therapeutics for this devastating form of cancer. Palmerolide A inhibits vacuolar ATPases, which are highly expressed in metastatic melanoma. Given the current level of understanding that macrolides often have microbial biosynthetic origins, that the holobiont metagenome has biosynthetic potential (26) and a diverse, yet persistent core microbiome is found in palmerolide A-containing *S. adareanum* (27) we have hypothesized that a microbe associated with *S. adareanum* is responsible for the biosynthesis of palmerolide A.

The core microbiome of the palmerolide A-producing ascidian *S. adareanum* in samples collected across the Anvers Island archipelago (n=63 samples; 27) is comprised of five bacterial phyla including Proteobacteria (dominating the microbiome), Bacteroidetes, Nitrospirae, Actinobacteria and Verrucomicrobia. A few candidate taxa in particular, were suggested to be likely palmerolide A producers based on relative abundance and biosynthetic potential determined by analysis of lineage-targeted biosynthetic capability (genera: *Microbulbifer, Pseudovibrio, Hoeflea*, and the family *Opitutaceae* (27). This motivated interrogation of the *S. adareanum* microbiome metagenome, with the goals of determining the metagenome-encoded biosynthetic potential, identifying candidate palmerolide A biosynthetic gene cluster (BGC)(s) and establishing the identity of the palmerolide A producing organism.

## Results and Discussion

### Identification of a candidate palmerolide A biosynthetic gene cluster

Microbial-enriched fractions of *S. adareanum* metagenomic DNA sequence from 454 and Ion Proton next generation sequencing (NGS) libraries (almost 18 billion bases in all) were assembled independently, then merged, resulting in ~ 145 Mbp of assembled bases distributed over 86,387 contigs (referred to as CoAssembly 1; supplemental material, Table S1). As the metagenome sequencing effort was focused on identifying potential BGCs encoding the machinery to synthesize palmerolide A, the initial steps of analysis specifically targeted those contigs in the assembly that were > 40 Kbp (102 contigs representing 0.12 % of all contigs, which represent 7.07 % of all reads mapped), as the size of the macrolide ring with 24 carbons would require a large number of polyketide modules to be encoded. This large fragment subset of CoAssembly 1 was submitted to antiSMASH v.3 (38) and more recently to v.5 (39). The results indicated a heterogeneous suite of BGCs, including a bacteriocin, two non-ribosomal peptide synthetases (NRPS), two hybrid NRPS -Type I PKS, two terpenes, and three *trans*-AT-PKS hybrid NRPS clusters (supplemental material, Table S2).

We predicted several functional characteristics of the BGC that would be required for palmerolide A biosynthesis which aided our analysis (see (40) for detailed retrobiosynthetic analysis). This included evidence of a hybrid nonribosomal peptide-polyketide pathway and enzymatic domains leading to placement of two distinct structural features of the polyketide backbone, a carbamoyl transferase that appends a carbamate group at C-11 of the macrolide ring, and an HMGCoA synthetase that inserts a methyl group on an acetate C-1 position of the macrolide structure (C-25). The antiSMASH results indicated that two of the three predicted hybrid NRPS *trans*-AT-Type I PKS contained the predicted markers. Manual alignment of these two contigs suggested near-identical overlapping sequence (36,638 bases) and, when joined, the merged contig resulted in a 74,672 Kbp BGC (Fig. 1C). The cluster size was in the range of other large *trans*-AT PKS encoding BGCs including pederin (54 Kbp; 41), leinamycin (135.6 Kbp; 42) as well as a *cis*-acting AT-PKS, jamaicamide (64.9 Kbp; 43). The combined contigs encompassed what appeared to be a complete BGC that was flanked at the start with a transposase and otherwise unlinked in the assembly to other contiguous DNA. The cluster lacked phylogenetically informative marker genes from which putative taxonomic assignment could be attributed.

The antiSMASH results suggested that the BGC appears to be novel with the highest degree of relatedness to pyxipyrrolone A and B (encoded in the *Pyxidicoccus* sp. MCy9557 genome (44), to which only 14% of the genes have a significant BLAST hit to genes in the metagenome-encoded cluster. The ketosynthase (KS) sequences (13 in all) fell into three different sequence groups (40). One was nearly identical (99% amino acid identity) to a previously reported sequence from a targeted KS study of *S. adareanum* microbiome metagenomic DNA (26). The other two were most homologous to KS sequences from *Allochromatium humboldtianum*, and *Dickeya dianthicola* in addition to a number of hypothetical proteins from environmental sequence data sets. The primary polyketide synthase (PKS) component of the BGC (labeled in Fig. 4C, Contig 9) includes 11 of the 13 KS domains which is consistent with a polyketide backbone which has 22 contiguous carbons since each elongation step adds a two-carbon unit. This further supports the hypothesis that this candidate BGC is likely responsible for palmerolide A biosynthesis. A detailed bioinformatic analysis into the step-wise biosynthetic mechanism was conducted by Avalon et al. (40).

### Taxonomic inference of palmerolide A BGC

Taxonomic attribution of the BGC was inferred using a real time PCR strategy targeting three coding regions of the putative palmerolide A BGC spanning the length of the cluster (acyltransferase, AT1; hydroxymethylglutaryl Co-A synthase, HCS, and the condensation domain of the non-ribosomal peptide synthase NRPS, Fig. 1C) to assay a *Synoicum* microbiome collection of 63 samples that have been taxonomically classified using Illumina SSU rRNA gene tag sequencing (27). The three gene targets were present in all samples ranging within and between sites at levels from ~ 7 × 10^1^ – 8 × 10^5^ copies per gram of host tissue (Fig. 2A). The three BGC gene targets co-varied across all samples (r^2^ > 0.7 for all pairs), with the NRPS gene copy levels slightly lower overall (mean: 0.66 and 0.59 copies per ng host tissue for NRPS:AT1 and NRPS:HCS respectively, n=63). We investigated the relationship between BGC gene copies per ng host tissue for each sample and palmerolide A levels determined for the same samples using mass spectrometry, however no correlation was found (R<0.03, n=63; 27). We then assessed the semi-quantitative relationship between the occurrence of SSU rRNA amplicon sequence variants (ASV, n=461; 27) and the abundance of the three palmerolide A BGC gene targets. Here, we found a robust correlation (R=0.83 – 0.99) between all 3 gene targets and a single amplicon sequence variant (ASV15) in the core microbiome (45). This ASV is affiliated with the *Opitutaceae* family of the Verrucomicrobium phylum. The *Opitutaceae* family ASV (SaM_ASV15) was a member of the core microbiome as it was detected in 59 of the 63 samples surveyed at varying levels of relative abundance and displayed strong correlations with the abundances of BGC gene targets (Fig. 2B, r^2^ =0.68 with AT1, 0.97 with HCS, and 0.69 with NRPS, n=63 for all). The only other correlations R > 0.5 were ASVs associated within the “variable” fraction of the microbiome, e.g., one low abundance ASV was present in 24 of 63 samples (45).

**Figure 2.**
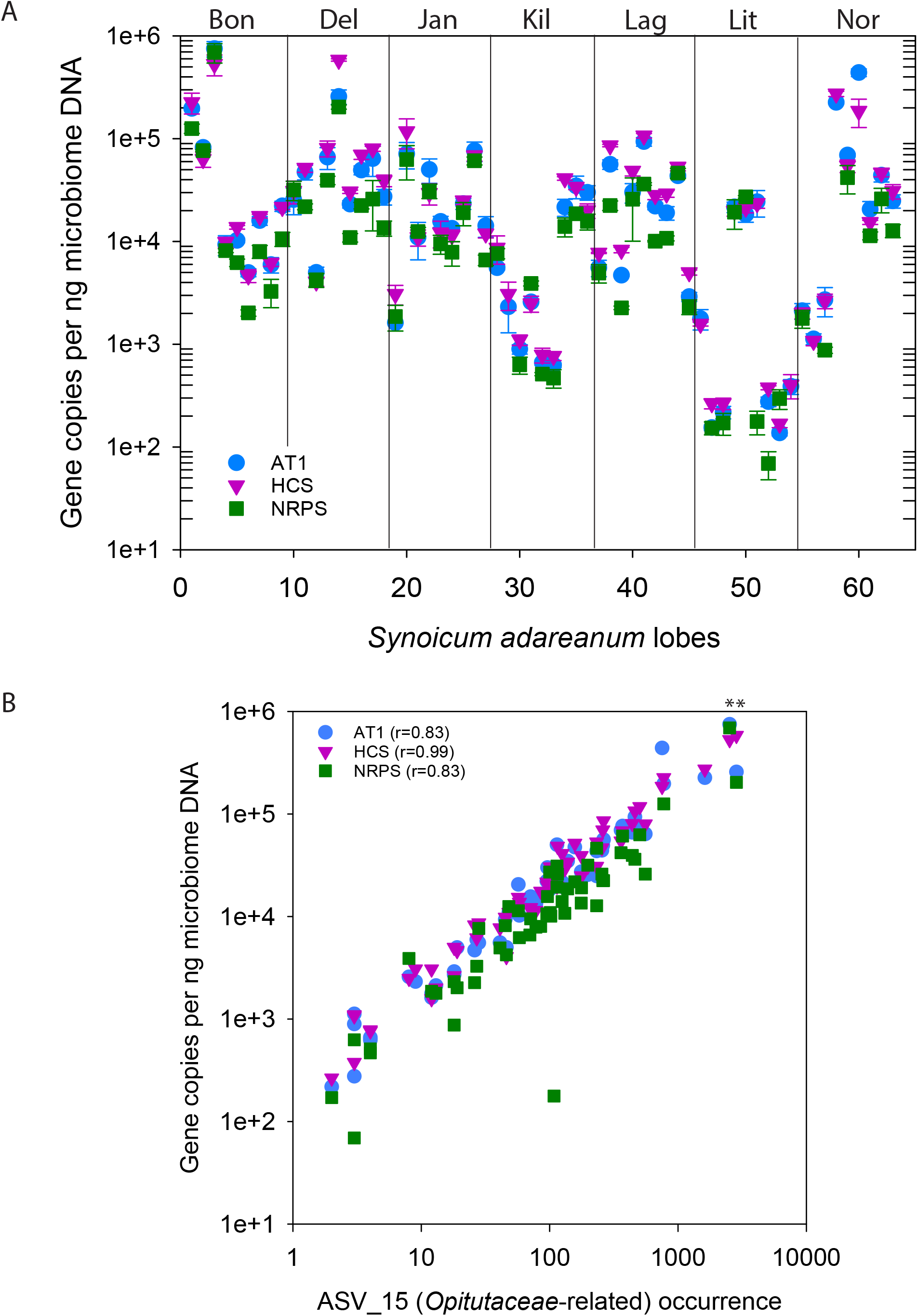
Abundances of real time PCR-targeted coding regions in the candidate pal biosynthetic gene cluster in Antarctic ascidian samples. (A) Gene copies estimated for three targeted coding regions (Acyltransferase, AT1; 3-hydroxy-methyl-glutaryl coenzyme A synthase, HCS; and the condensation domain of a non-ribosomal peptide synthase; NRPS) in the candidate *pal* biosynthetic gene cluster surveyed over 63 DNA extracts derived from microbial cell preparations enriched from the Antarctic ascidian *Synoicum adarenum*. Nine samples were collected at each of seven sites: Bon, Bonaparte Point; Del, Delaca Island; Jan, Janus Island, Kil, Killer Whale Rocks; Lag, Laggard Island, Lit, Litchfield Island; Nor, Norsel Point (27). (B) Relationship between gene copy number for the three gene targets and the 16S rRNA gene ASV occurrences of Opitutaceae-related ASV_15 across a 63 *S. adareanum* microbial DNA sample set. * indicates samples Bon-1C-2011 and Del-2b-2011 that were selected for PacBio sequencing.

This result supports the finding of Murray et al. (27) in which gene abundance and natural product chemistry do not reflect a 1:1 ratio in this host-associated system. Neither the semi-quantitative measure of ASV copies nor the real time PCR abundance estimates of the three biosynthetic gene targets correlated with the mass-normalized levels of palmerolide A present in the same samples. As discussed (27), this is likely a result of bioaccumulation in the ascidian tissues. This result provided strong support that the genetic capacity for palmerolide A production was associated with a novel member of the *Opitutaceae*, a taxonomic family with representatives found across diverse host-associated and free-living ecosystems. Although the biosynthetic capacity of this family is not well known (46), recent evidence (36) suggests this family may be a fruitful target for cultivation efforts and natural product surveys.

### Assembly of the palmerolide BGC-associated *Opitutaceae*-related metagenome assembled genome (MAG)

With metagenomes some genomes come together easily – while others present compelling puzzles to solve. Assembly of the *pal* BGC-containing *Opitutaceae* genome was the result of a dedicated effort of binning contigs, gene searches, additional sequencing of samples with high BGC titer, and manual, targeted assembly. Binning efforts with CoAssembly 1 did not result in association of the *pal* BGC with an associated metagenome assembled genome (supplemental materials and methods, Table S3). Therefore, a further round of metagenome sequencing using long read technology (Pacific Biosciences Sequel Systems technology; PacBio) ensued.

The 16S rRNA gene ASV occurrence (27) and real time PCR data were used to guide *S. adareanum* sample selection for sequencing. Two ascidian samples (Bon-1C-2011 and Del-2B-2011) with high *Opitutaceae* ASV occurrences (ASV_015; > 1000 sequences each – relative abundance of ~ 13.3-15.3 % compared to an overall average of 1.3 ± 2.77% across the 63 samples respectively, (27) and high BGC gene target levels (i.e., Bon-1C-2011 (copies per ng ± s.d., n=3): 6.93 ± 1.48 × 10^5^ NRPS; 7.52 ± 1.29 × 10^5^ AT1; 5.32 ± 1.22 × 10^5^ HCS and Del 2B-2011 (copies per ng ± s.d., n=3): 2.04 ± 0.10 × 10^5^ NRPS; 2.57 ± 0.41 × 10^5^ AT1; 5.84 ± 0.16 × 10^5^ HCS) were selected for PacBio sequencing. This effort generated 28 GB of data that was used to create a new hybrid CoAssembly 2 which combined all three sequencing technologies. Similar to the assembly with the *Mycale hentscheli*-associated polyketide producers (47), the long-read data set improved the assembly metrics, and subsequent binning resulted in a highly resolved *Opitutaceae*-classified bin (45, supplemental material, Table S3). Interestingly however, the palmerolide BGC contigs still did not cluster with this bin, which we later attributed to binning reliance on sequence depth.

We used PacBio circular consensus sequence (CCS) reads to generate and manually edit the assembly for our *Opitutaceae* genome of interest. The resulting 4.3 Mbp genome (Fig. 3A) had a GC content of 58.7% and was resolved into a total of 10 contigs. Five of the contigs (Contigs 1-5) were unique while contigs 6-10 represented highly similar repeated units of the *pal* BGC (labeled *pal* BGC 1, 2, 3, 4, and 5) beginning and ending in linkage gaps. Support for the assembly of a near-complete genome includes manual analysis of the ends of the five unique contigs and underlying read data linking each end of all five contigs to parts of the repeated contigs. The structure of the five repeated contigs was manually evaluated (Fig.1) and while each copy had unique features, all were found to harbor portions of the palmerolide BGC. While the exact placement of each of the five palmerolide contigs could not be positioned with respect to the five unique contigs, the data support a single circular chromosome. Analyses further supporting a complete genome includes the presence of a complete ribosomal operon and estimated CheckM completeness of 96.04 % based on marker genes (presence of which were identified using MetaERG annotation of the *Opitutaceae* genome, Table S3), and 45 tRNA genes.

**Figure 3.**
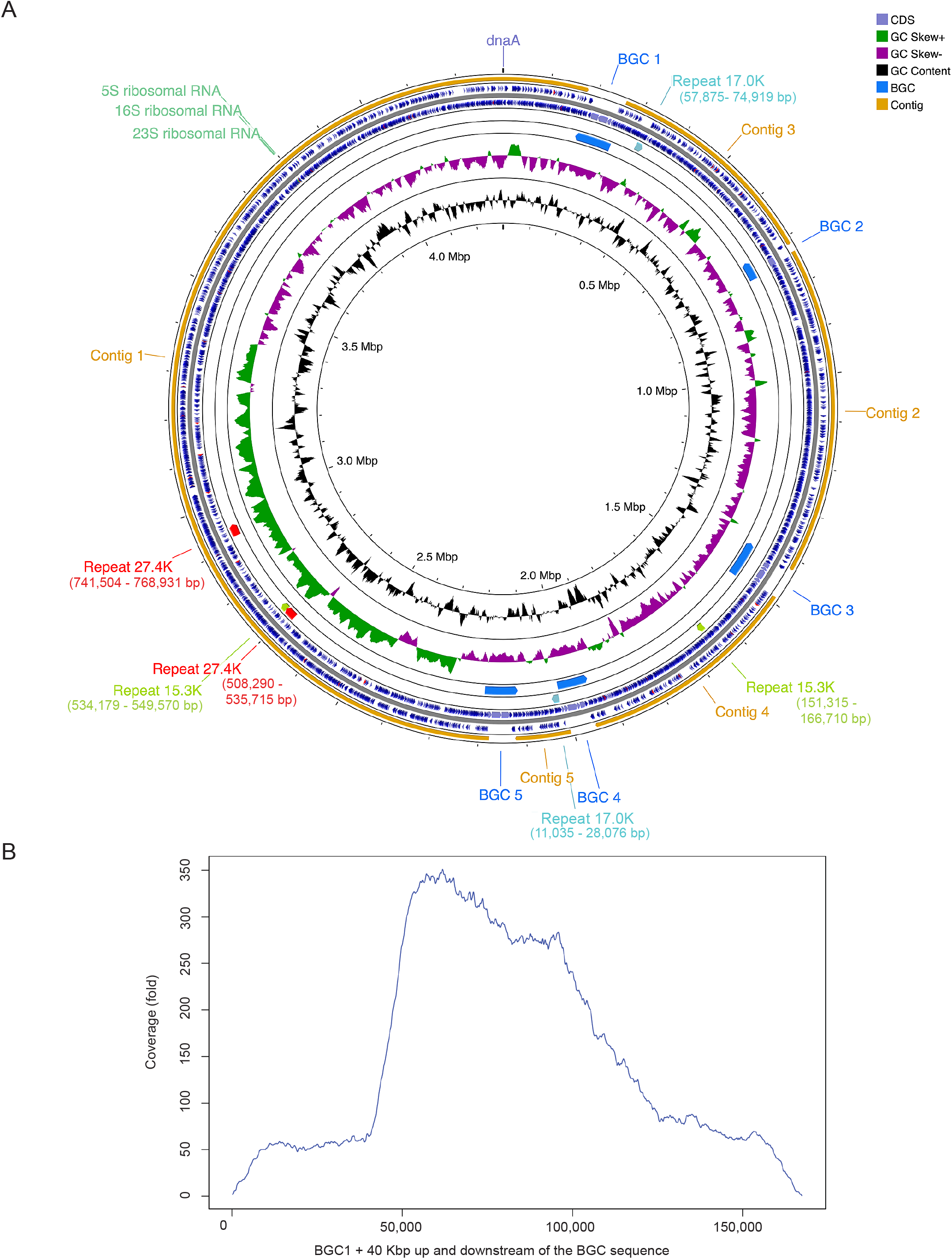
Genome maps of assembled MAG, *Candidatus* Synoicohabitans palmerolidicus and evidence of multi-copy biosynthetic gene clusters. (a) The 4,297,084 bp gene map is oriented to dnaA at the origin. One possible assembly scenario of the *Ca*. S. palmerolidicus genome is shown as the order of the contigs and palmerolide BGC’s are not currently known. In addition to the 5 BGCs, three other internally repetitive regions were identified (15.3, 17.0 and 27.4 Kbp). The genes and orientation are shown with blue and tRNA indicated in red. (b) To demonstrate depth of coverage outside and inside the BGC regions, CCS reads from sample Bon-1C-2011 and Del-2b-2011 were mapped to a 167.6 Kbp region. The profile extends 40 Kbp into the genome on either side of the BGC where depth of coverage averages 60 fold, while in the BGC depth of coverage varies across the BGC given differences in cover across the BGC, the highest cover is 5X, or ~ 300 fold, supporting the finding of 5 repeats encoding the BGC.

Alignment of all five repeated contigs to the longest palmerolide-containing BGC revealed a long (36,198 base) repeated region that was shared between all 5 contigs with some substantial differences at the beginning of the cluster and only minor differences at the end, indicating 3 full length, and 2 shorter palmerolide BGC-containing contigs (Fig. 1 and 3). This was consistent with coverage estimates based on read-mapping that suggested lower depth at the beginning of the cluster (Fig. 3B). BGC 1 and 3 are nearly identical (over 86,135 bases) with only 2 single nucleotide polymorphisms (SNPs) and an additional 1,468 bases in BGC1 (237 bases at the 5’ end and 1231 bases at the 3’ end). BGC 4 is 13,470 bases shorter than BGC1 at the 5’ end, and 5 bases longer than BGC 1 at the 3’ end. Alignment of the real time PCR gene targets to the 5 *pal* BGCs provided independent support for the different lengths of the 5 BGCs, as the region targeted by the NRP primers was missing in two of the *pal* BGCs, thus explaining lower NRP: AT or NRP:HCS gene dosages reported above.

Interestingly, precedent for naturally occurring multi-copy BGCs to our knowledge, has only recently been found in one other bacterium, an ascidian (*Lissoclinum* sp.)-associated *Opitutaceae, Candidatus* Didemnitutus mandela, linked to cytotoxic mandelalide polyketides (36). Likewise, we can invoke a rationale similar to (36) that multiple gene clusters may be linked to biosynthesis of different palmerolide derivatives; see Avalon et al. (40) for retro-biosynthetic predictions of these clusters and annotation of putative enzymatic functions. Gene duplication, loss, and rearrangement processes over evolutionary time, likely explain the source of the multiple copies. At present we do not yet understand the regulatory controls, whether all five are actively transcribed, in situ function and how this may vary amongst host microbiomes.

### Phylogenomic characterization of the Opitutaceae-related MAG

The taxonomic relationship of the *Opitutacae* MAG to other Verrucomicrobiota was assessed using distance-based analyses with 16S rRNA and average amino acid identity (AAI). Then, it was classified using the GTDB-Tk tool (48), and a phylogenomic analysis based on concatenated ribosomal protein markers. Comparison of 16S rRNA gene sequences amongst other Verrucomicrobia with available genome sequences (that also have 16S rRNA genes; supplemental material, Fig. S1) suggests that the nearest relatives are *Cephaloticoccus primus* CAG34 (similarity of 0.9138), *Optitutus terrae* PB90-1 (similarity of 0.9132) and *Geminisphaera coliterminitum* TAV2 (similarity of 0.9108). The *Opitutaceae*-affiliated MAG sequence is identical to a sequence (uncultured bacterium clone Tun-3b A3) reported from the same host (*S. adareanum*) in a 2008 study (26); bootstrapping supported a deep branching position in the *Opitutaceae* family. Thus, this Opitutaceae-related MAG appears to be unique – its 16S rRNA gene sequence was not found in bacterioplankton amplicon surveys from the Anvers Island archipelago (n=604 amplicon sequences) and a culture collection reported previously (27), nor were any sequences with identities higher than 95% found following against a large bacterioplankton amplicon data set from further north in the Antarctic Peninsula (32,941 sequence clusters derived from 44.6 million sequences; NCBI Bioproject PRJNA316748). Likewise, when searched against the global nr GenBank database representing environmental data sets the highest sequence identities found so far are < 96% identity over 88% of the complete sequence.

When characterizing the MAG using AAI metrics (average nucleotide identity, ANI, found no closely related genomes) the closest genomes were environmental metagenome assemblies from the South Atlantic TOBG_SAT_155 (53.08 % AAI) and WB6_3A_236 (52.71 % AAI); and the two closest isolate type genomes were *Nibricoccus aquaticus* str. NZ CP023344 (52.82 % AAI) and *Opitutus terrae* str. PB90 (52.75 % AAI). The Microbial Genome Atlas (MiGA) support for the MAG belonging in the *Opitutaceae* family was weak (p-values of 0.5). Attempts to classify this MAG using GTDB-Tk (47) were hampered by the fact we have no real representative in the genome databases, resulting in low confidence predictions at the species or genus levels (see the supplemental material for details).

Verrucomicrobia exhibit free-living and host-associated lifestyles in a multitude of terrestrial and marine habitats on Earth. We performed a meta-analysis of Verrucomicrobia genomes, with an emphasis on marine and host-associated *Opitutaceae*, to establish more confidence in the phylogenetic position of the *Opitutaceae* MAG. The analysis was based on 24 conserved proteins – 21 ribosomal proteins and three additional conserved proteins (InfB, lepA, pheS). The diversity of the *Opitutaceae* family, and of Verrucomicrobia in general, is largely known from uncultivated organisms in which there are 20 genera in GTDB (release 05-RS95), 2 additional genera in the NCBI taxonomy database, and numerous unclassified single amplified genomes (SAGs); in all, only eight genera have cultivated representatives. Given the uneven representations of the 24 proteins across all (115) genomes assessed (MAGs and SAGs are often incomplete), we selected a balance of 16 proteins across 48 genomes to assess phylogenomic relatedness across the *Opitutaceae* (Fig. 4). Here too, as seen with the 16S rRNA gene phylogenetic tree, the *S. adareanum*-*Opitutaceae* MAG held a basal position compared to the other *Opitutaceae* genomes in the analysis.

**Figure. 4.**
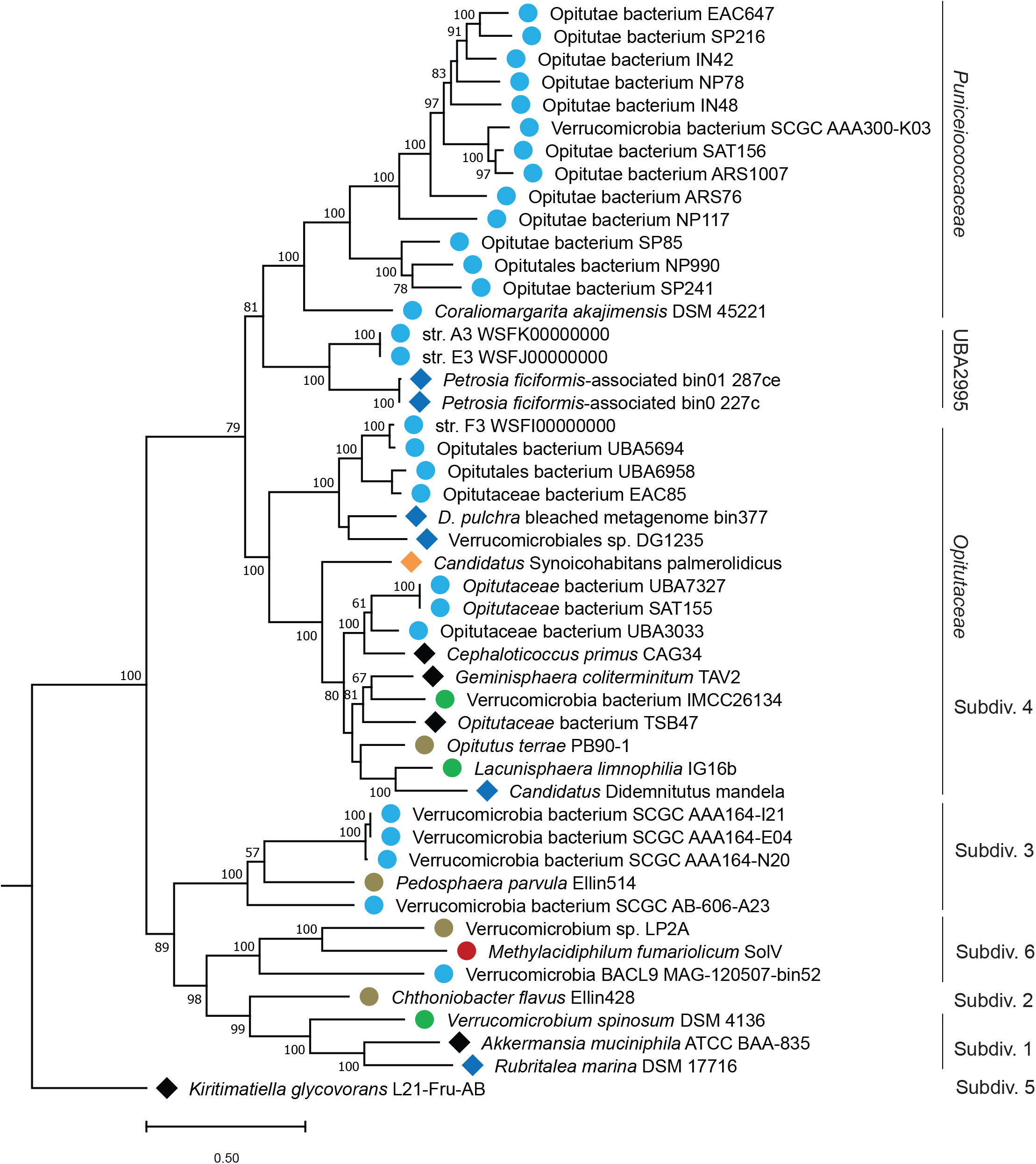
Maximum likelihood phylogenomic tree showing 48 verrucomicrobia genomes. Phylogenomic relationship of *Candidatus* Synoicohabitans palmerolidicus (*Opitutaceae* bin 8) with respect to other mostly marine, and host-associated verrrucomicrobia subdivision 4 and other genomes. The tree is based on 16 concatenated ribosomal proteins (5325 amino acids) common across 48 Verrucomicrobia genomes. Distance was estimated with RAxML with 300 bootstrap replicates. Symbols designate environmental origins of the organisms: free-living are represented by circles: light blue - marine, green– freshwater, red – hydrothermal mud, brown – soils. Host-associated taxa from marine systems with blue diamonds, and from terrestrial systems with black diamonds.

### *Ca*. S. palmerolidicus relative abundance estimates and ecological inference

The relative abundance of *Opitutaceae* bin 8 was estimated in the shotgun metagenomic samples by mapping the NGS reads back to the assembled MAG across the four *S. adareanum* samples collected. This indicated varying levels of genome coverage in the natural samples, with the two samples selected based on real time PCR-quantified high BGC copy number being clearly enriched in this strain (44.70 % of reads mapped to Bon-1C-2011 and 36.78 % to Del-2B-2011, Table 1). These levels are higher than estimates of relative abundance derived from the 16S rRNA gene amplicon surveys (estimated at 13.33 and 15.34 % respectively) for the same samples. This is likely a result of the single-copy nature of the ribosomal operon in *Opitutaceae* bin 8 vs. other taxa with multiple rRNA operon copies that could thus be over-represented in the core microbiome library (e.g., *Pseudovibrio* sp. str. PSC04-5.I4 has 9 and *Microbulbifer* sp. is estimated at 4.1 ± 0.8 based on 9 finished *Microbulbifer* genomes available at the Integrated Microbial Genomes Database). All host *S. adareanum* lobes surveyed (n=63) in the Anvers Island regional survey contained high levels (0.49 – 4.06 mg palmerolide A x g^-1^ host dry weight) of palmerolide A (27), and variable, yet highly concordant levels of the *pal* BGCs and 16S rRNA ASV levels (Fig. 2). Despite the natural population structure sampled here (four single host lobes), the bin-level sequence variation was low (ranging from 72-243 SNPs) when the PacBio reads were mapped back to the Opitutaceae bin 8 (Table 1). This suggests maintenance of a relatively invariant population at the spatial and temporal scales of this coastal Antarctic region while highlighting our limited understanding of the biogeographical extent of the *S. adareanum*-symbiont-palmerolide relationship across a larger region of the Southern Ocean.

**Table 1.**
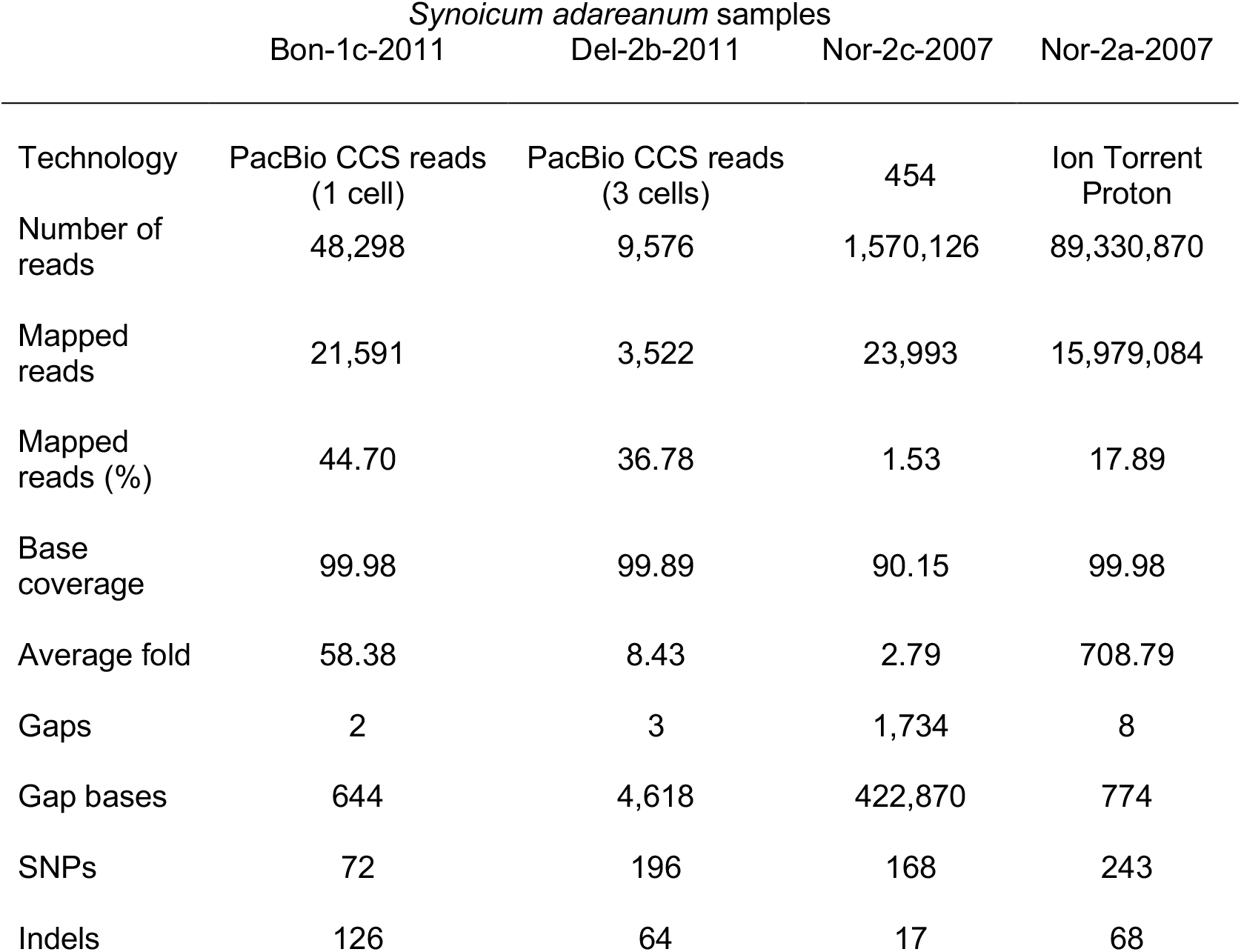
Metagenomic reads from 4 different samples were mapped back to the *Ca*. S. palmerolidicus MAG.

|  | *Synoicum adareanum* samples |  |  |  |
| --- | --- | --- | --- | --- |
|  | Bon-1c-2011 | Del-2b-2011 | Nor-2c-2007 | Nor-2a-2007 |
| Technology | PacBio CCS reads (1 cell) | PacBio CCS reads (3 cells) | 454 | Ion Torrent Proton |
| Number of reads | 48,298 | 9,576 | 1,570,126 | 89,330,870 |
| Mapped reads | 21,591 | 3,522 | 23,993 | 15,979,084 |
| Mapped reads (%) | 44.70 | 36.78 | 1.53 | 17.89 |
| Base coverage | 99.98 | 99.89 | 90.15 | 99.98 |
| Average fold | 58.38 | 8.43 | 2.79 | 708.79 |
| Gaps | 2 | 3 | 1,734 | 8 |
| Gap bases | 644 | 4,618 | 422,870 | 774 |
| SNPs | 72 | 196 | 168 | 243 |
| Indels | 126 | 64 | 17 | 68 |

Several questions remain with regard to the in situ function of palmerolide A (a eukaryotic V-ATPase inhibitor in human cell line assays (28) in this cryohabitat: how and why is it bioaccumulated by the host? Overall, the study of natural products in high latitude marine ecosystems is in its infancy. This palmerolide producing, ascidian-associated, *Opitutaceae* provides the first Antarctic example in which a well-characterized natural product has been linked to the genetic information responsible for its biosynthesis. Gaining an understanding of environmental and biosynthetic regulatory controls, establishing integrated transcriptomic, proteomic, and secondary metabolome expression in the environment will also reveal whether the different clusters are expressed in situ. In addition to ecological pursuits, the path to clinical studies of palmerolide will require genetic or cultivation efforts. At present, we hypothesize that cultivation of *Opitutaceae* bin 8 may be possible, given the lack of genome reduction or of other direct evidence for host-associated dependencies (e.g., a number of central carbohydrate and energy metabolism pathways appear to be present).

### *Candidatus* Synoicihabitans palmerolidicus genome attributes

The Antarctic ascidian, *Synoicum adareanum*, harbors a dense community of bacteria that has a conserved core set of taxa (27). The near complete ~ 4.30 Mbp *Opitutaceae* bin 8 metagenome assembled genome (Fig. 3) represents one of the core members. This MAG is remarkable in that it encodes for five 36-74 Kbp copies of the candidate BGCs that are implicated in biosynthesis of palmerolide A and possibly other palmerolide compounds. Intriguingly, this genome does not seem to show evidence of genome reduction as found in *Candidatus* Didemnitutus mandela (36); the other ascidian-associated *Opitutaceae* genome currently known to encode multiple BGC gene copies. This is the first *Opitutaceae* genome characterized from a permanently cold, ~ -1.8 - 2 °C, often ice-covered ocean ecosystem. This genome encodes one rRNA operon, 45 tRNA genes, and an estimated 5058 coding sequences. Based on the low (< 92%) SSU rRNA gene identity and low (< 54% AAI) values to other genera in the *Opitutaceae*, along with the phylogenomic position of the *Opitutaceae* bin 8, the provisional name “*Candidatus* Synoicihabitans palmerolidicus” (*Ca*. S. palmerolidicus) is proposed for this novel verrucomicrobium. The genus name *Synoicihabitans* (Syn.o.i.ci.ha’bitans. N.L. neut. N. *Synoicum* a genus of ascidians; L. pres. part *habitans* inhabiting; N.L. masc. n.) references this organism as an inhabitant of the ascidian genus *Synoicum*. The species name *palmerolidicus* (<u>pal.me.ro.li</u>’di.cus. N.L. neut. n. *palmerolidum* palmerolide; N.L. masc. adj.) designates the species as pertaining to palmerolide.

The GC content of 58.7% is rather high compared to other marine *Opitutaceae* genomes (ave. 51.49 s.d. 0.02, n=12), yet is ~ average for the family overall (61.58 s.d. 0.06, n=69; supplemental material, Table S4). MetaERG includes metagenome assembled genomes available in the GTDB as a resource for its custom GenomeDB that new genomes are annotated against. This was a clear advantage in annotating the *Ca*. S. palmerolidicus genome as Verrucomicrobia genomes are widely represented by uncultivated taxa. Likewise, antiSMASH was an invaluable tool for *pal* BGC identification and domain structure annotation. This formed the basis to derive a predicted step-wise mechanism of *pal* biosynthesis (40).

### *Ca*. S. palmerolidicus genome structure, function and host-associated features

Beyond the *pal* BGCs, the *Ca*. S. palmerolidicus genome encodes a variety of additional interesting structural and functional features that provide insight its lifestyle. Here we provide but a brief synopsis. In addition to the repeated BGCs, three additional repeated elements with two nearly identical copies each (15.3 Kbp, 17.0 Kbp, 27.4 Kbp) were identified during the assembly process (Fig. 3A). These coded for 20, 25 and 41 CDSs respectively, were in some cases flanked by transposase/integrases (both internal and proximal) and had widespread homology with Verrucomicrobia orthologs. The contents of the three repetitive elements were not shared amongst each other.

Annotations were assigned to a little more than half of the CDSs in the 15.3 Mbp repeat which is predicted to encode for xylose transport, two sulfatases, two endonucleases and a MacB-like (potential macrolide export) periplasmic core protein. Xylan might be sourced from seaweeds (49) or even the ascidian, as it is a minor component of the tunic cellulose (50). Related to this, an endo-1,4-beta-xylanase which has exoenzyme activity in some microorganisms (51) was identified elsewhere in the genome suggesting the potential for xylose metabolism. Altogether, eight sulfatase copies were identified in this host-associated genome (four in the 15.5 Mbp repeat elements). These may be involved in catabolic activities of sulfonated polysaccharides, and possibly as *trans*-acting elements in palmerolide biosynthesis (40). In addition to the MacB-like CDSs found in this repeat, 13 other MacB-homologs were present in the genome – none of which were associated with the *pal* BGCs (supplemental material, Fig. S2a). MacB is a primary component of the macrolide tripartite efflux pump that operates as a mechanotransmission system involved both in antibiotic resistance and antibiotic export depending on the size of the macrolide molecule (52). However, two additional elements required for this pump to be functional, an intramembrane MacA and an outer membrane protein TolC, were not co-located elsewhere in the genome. MacA may be missing, as searches against the of *Ca*. S. palmerolidicus genome with to two other verrucomicrobia-associated MacA CDSs were not identified using BLAST queries (Peat Soil MAG SbV1 SBV1_730043 and *Ca*. Udaeobacter copiosis KAF5408997.1; (53). At least nine MacB CDSs were flanked by a FstX-like permease family protein; the genomic structures of which were quite complex including several with multiple repeated domains. Detailed transporter modeling is beyond the scope of this work, but it is likely that these proteins are involved in signaling of cell division machinery rather than macrolide transport (54).

Predicted CDSs in the 17.0 Mbp repeat included sugar binding and transport domains, as well as domains encoding rhamnosidase, arabinofuranosidase, and other carbohydrate catabolism functions. About half the proteins encoded in the 27.4 Mbp repeat were unknown in function, and of the remaining characterized suggested diverse potential functional capacities. For example, a zinc carboxypeptidase (1 of 3 in the genome), multidrug and toxic compound transporter (MatE/NorM), and an exodeoxyribonuclease were identified.

The *Ca*. S. palmerolidicus genome has a number of features that suggest it is adapted to a host-associated lifestyle, and several of these features were reported recently for two related sponge-associated Opitutales metagenome bins (*Petrosia ficiformis-*associated bins 0 and 01, Fig. 4; (55). These include identification of a bacterial microcompartment (BMC) ‘super locus’. Such loci were recently reported to be enriched in host-associated Opitutales genomes when compared to free-living relatives. The structural proteins for the BMC were present as were other conserved Planctomyces-Verrucomicrobia BMC genes (56). As in the sponge *Pectoria ficiformis* metagenome bins, enzymes for carbohydrate (rhamnose) catabolism and modification were found adjacent to the BMC locus (supplemental material, Fig. S3), in addition to the two that were found in the 27.4 Mbp repeat. The genome did not appear to encode the full complement of enzymes required for fucose metabolism, though a few alpha-L-fucosidases were identified. Further evidence for carbohydrate metabolism was supported through genome similarity searches with the CAZY database (57), including 7 identified carbohydrate binding modules, a carbohydrate esterase, 14 glycoside hydrolases, 6 glycosyl transferases and a polysaccharide lyase. In addition, three bacterial cellulases (PF00150, cellulase family A; glycosyl hydrolase family 5) were identified, all with the canonical conserved glutamic acid residue. These appear to have different evolutionary histories in which each variant has nearest neighbors in different bacterial phyla (supplemental material, Fig. S2b) matching between 68% identity for Protein J6386 03765 to *Lacunisphaera limnophila*, 57.5% identity for Protein J6386 22340 with a cellulase from a shipworm symbiont *Alteromonadaceae* (*Terridinibacter* sp.), and 37.5% sequence identity to a Bacteroidetes bacterium. This suggests the potential for cellulose degradation – which is consistent with ascidians being the only animals known to produce cellulose for its skeletal structure (50). In addition to the BGCs, the enzymatic resources in this genome (e.g., xylan and cellulose hydrolysis) are a treasure trove rich with biotechnological potential. Support for a Type II secretion system (i.e. GspD, GspE, GspG and GspO), common to gram-negative bacteria which secrete folded proteins (e.g. hydrolytic enzymes required for survival in host environments) from the periplasm into the extracellular environment (58) were detected in the *Ca*. S. palmerolidicus genome.

Chemotaxis and flagellar biosynthesis are factors required for horizontally acquired symbionts, as has been established with the *Vibrio fischeri-Euprymna scolopes* symbiosis (59). The *Ca*. S. palmerolidicus genome encodes a chemotaxis system (e.g. CheA, CheW, CheR, CheB, Che C, methyl-accepting chemotaxis domains) with flagellar motors in addition to a number of other elements of flagellar biosynthesis, which is consistent with horizontal acquisition by the host. The methyl-accepting chemotaxis system in *V. fischeri* responds to chitobiose production by *E. scolopes* (59) – it will be interesting to further unravel the details of Ca. S. palmerolidicus’ cellular biology to understand the chemotaxis stimulants, preferential cellular localization of palmerolide production and resistance mechanisms of the host to the potent vacuolar ATPase, as well as products made by others in the *S. adareanum* microbiome.

Other indicators of host-association and palmerolide production include T-A domains, multidrug exporters, and the potential for palmerolide transport and cofactor biosynthesis. T-A domains were also prevalent in the *Petrosia ficiformis-*associated bins 0 and 01 (55). The *Ca*. S. palmerolidicus genome encoded at least 22 TA-related genes including multiple MazG and AbiEii toxin type IV TA systems, AbiEii-Phd_YefM type II toxin-antitoxin systems, along with genes coding for PIN domains, Zeta toxin, RelB, HipA, MazE and MraZ. In addition to the MatE (found in the 27.4 Mbp repeat) two other multidrug export systems with homology to MexB and MdtB were identified. This analysis also resulted in identifying a putative AbiEii toxin (PF13304) with homology to SyrD, a cyclic peptide ABC type transporter that was present in all 5 BGCs (Fig. 1C; with 52.7% BLAST percent identity to a *Desulfamplus* sp. homolog over the full length of the protein, and a variety of other bacteria including an *Opitutaceae*-related strain with similar levels of identity). This transporter is encoded downstream of the large polyketide gene clusters following the acyl transferase domains and precedes the predicted *trans*-acting domains at the 3’ end of the BGC. Given its genomic position, this protein is a candidate for palmerolide transport. The *Ca*. S. palmerolidicus genome also encodes the potential for pantothenate biosynthesis via ilvD, ilvE, panB, ilvC, panD, and panC (60), which is consistent with palmerolide biosynthesis in which the 4’-phosphopantetheinyl prosthetic group interacts with acyl carrier proteins for multimodular assembly. Some symbionts, e.g. *Ca*. Entotheonella sp., are auxotrophic for pantothenate and likely acquire it from other microbiome members (61).

Unlike in *Ca*. D. mandela (36), there does not appear to be ongoing genome reduction, which may suggest that the *S. adareanum*-*Ca*. S palmerolidicus relationship is more recent, and/or that the relationship is commensal rather than interdependent. Likewise, we suspect that the pseudogene content may be high as several CDS appear to be truncated, in which redundant CDS of varying lengths were found in several cases (including the MacB). There is evidence of lateral gene transfer acquisitions of cellulase and numerous other enzymes that may confer ecological advantages through the evolution of this genome. Similarly, the origin of the *pal* BGCs and how recombination events play out in the success of this Antarctic host-associated system in terms of adaptive evolution (62), not to mention the ecology of *S. adareanum* is a curiosity. This phylum promises to be an interesting target for further culture-based and cultivation-free studies – particularly in the marine environment, in which they have been less well studied compared to terrestrial free-living and host associated systems.

Together, it appears that the genome of *Ca*. S. palmerolidicus is equipped for life in this host-associated interactive ecosystem that stands to be one of the first high latitude marine invertebrate-associated microbiomes with a genome-level understanding – and one that produces a highly potent natural product, palmerolide A. This system holds promise for future research now that we have identified the producing organism and *pal* BGC. We still have much to learn about the ecological role of palmerolide A – if is it involved predation avoidance, antifouling, antimicrobial defense or some other yet to be recognized aspect of life in the frigid, often ice-covered and seasonally light-limited waters of the Southern Ocean.

## Materials and Methods

### Sample Collection

*S. adareanum* lobes were collected in the coastal waters off Anvers Island, Antarctica and stored at -80 °C until processing (supplemental material, Table S1). See the supplemental material and (27) for details of sample collection, microbial cell preparation and DNA extraction.

### Metagenome sequencing

Three rounds of metagenome sequencing were conducted, the details of which are in the supplemental material. This included an initial 454 pyrosequencing effort with a bacterial-enriched metagenomic DNA preparation from *S. adareanum* lobe (Nor2c-2007). Next, an Ion Proton System was used to sequence a metagenomic DNA sample prepared from *S. adareanum* lobe Nor2a-2007. Then two additional *S. adareanum* metagenome DNA samples (Bon-1C-2011 and Del-2b-2011) selected based on high copy numbers of the palmerolide A BGC (see real time PCR Methods, supplemental material) and sequenced using Pacific Biosciences Sequel Systems technology.

### Metagenome assembly, annotation and binning

Raw 454 metagenomic reads (1,570,137 single end reads, 904,455,285 bases) were assembled by Newbler (63) v2.9 (Life Technologies, Carlsbad, CA, flags: -large -rip -mi 98 -ml 80), while Ion Proton metagenomic reads (89,330,870 reads, 17,053,251,055 bases) were assembled using SPAdes (64) v3.5 (flags: --iontorrent). Both assembled datasets were merged with MeGAMerge (65) v1.2 and produced 86,387 contigs with a maximum contig size 153,680 and total contig size 144,953,904 bases (CoAssembly 1). To achieve more complete metagenome coverage and facilitate metagenome assembled genome assembly, a Circular Consensus Sequence (CCS) protocol (PacBio) was used to obtain high quality long reads on two samples Bon-1C-2011 and Del-2b-2011. The 5,514,426 PacBio reads were assembled with aforementioned assembled contigs (CoAssembly 1) on EDGE Bioinformatics using wtdbg2 (66), a fast and accurate long-read assembler. The contigs were polished with three rounds of polishing by Racon (67) into a second Coassembly (CoAssembly 2) which has 4,215 contigs with a maximum contig size 2,235,039 and total size 97,970,181 bases. Lastly, a manual approach was implemented to arrive at assembly of the MAG of interest the details of which are described in the supplemental material.

The contigs from both co-assemblies 1 and 2 were submitted initially to the EDGE bioinformatics platform (68) for sequence annotation using Prokka (69) v1.13 and taxonomy classification using BWA (70) mapping to NCBI RefSeq (Version: NCBI 2017 Oct 3). Bioinformatic predictions of natural product potential was performed using the antibiotics and secondary metabolite analysis shell (antiSMASH, bacterial versions 3.0, 4.0 and 5.0 (38, 39, 71). This tool executed contig identification, annotation and analysis of secondary metabolite biosynthesis gene clusters on both CoAssemblies 1 and 2 (> 1 Kbp and > 40 Kbp data sets). As most of our attention was focused on analysis the *Ca*. S. palmerolidicus assembled metagenome, we also used MetaERG (72) as the primary pipeline for metagenome annotation of the ten final contigs in addition to NCBI’s PGAP pipeline (see (45) for GFF dataset). There were 5186 coding sequences predicted in the MetaERG annotation and 5186 in NCBI’s PGAP annotation.

MaxBin (73) and MaxBin2 (74) were used to form metagenome bins for both CoAssembly 1 and 2. CheckM v1.1.11 (75) and v.1.1.12, and GTDB-Tk v.1.0.2 (48) were used to verify bin quality and taxonomic classification. See supplemental material for details. In order to assess the representation of assembled *Opitutaceae* genome across the 4 environmental samples used for metagenome sequencing (resulting from MaxBin2 binning of CoAssembly 2), we used BWA to map the CCS reads to each metagenome data set.

### Real time PCR

Gene targets (non-ribosomal peptide synthase, acyltransferase, and 3-hydroxymethylglutaryl coenzyme A synthase) were selected at different positions along the length of the candidate BGC. Supplemental material, Table S5 lists the primer and the GBlocks synthetic positive control sequence. Metagenomic DNA extracts from a large *S. adareanum* sample set (n=63 *S. adareanum* lobes from 21 colonies), all containing high levels of palmerolide A (27), were screened with the real time PCR assays on a Quant Studio 3 (Thermo Fisher Scientific, Inc.; see supplemental material text for details of controls and analysis).

### Phylogenomic analyses

A phylogenomic analysis of the assembled *Opitutaceae* MAG was conducted based on shared ribosomal RNA and ribosomal proteins amongst 46 and 48 reference genomes respectively, out of 115 genomes in total, mined from various databases (NCBI, GTDB and IMG) for uncultivated and cultivated microorganisms identified in the Verrucomicrobia phylum (supplemental material, Table S6). The details of these analyses are described in the supplemental material. In addition, we used MiGA (NCBI Prokaryotic taxonomy and the environmental TARA Oceans (Tully) databases; accessed August 2020) and GTDB-Tk (ver. 1.3.0) tools for MAG taxonomic classification.

Phylogenetic analysis of the MacB CDS sequences were retrieved from MetatERG annotated *Ca*. S. palmerolidicus contigs and homologs were retrieved from the NCBI based on BLAST results. Maximum likelihood analysis was conducted on 994 aligned (MUSCLE) positions using RAxML v.8.2.12 using the PROTGAMMALG model and 550 bootstrap replicates. For the phylogenetic analysis of the cellulase CDS, homologs were retrieved from the NCBI based on BLAST results resulting in 19 sequences and 496 aligned positions (ClustalOmega) was also conducted using RAxML v.8.2.12 under the PROTGAMMALG model of evolution with 1000 bootstraps.

## Supporting information

All Supplemental Material

## Data Availability

The data (biosamples, SRA depositions) associated with this study is associated with NCBI BioProject PRJNA662631. The *Ca*. S. palmerolidicus biosample ID is SAMN18473105 and genome accession JAGGDC000000000. An annotation data set (.gff) for the Ca. S. palmerolidicus MAG resulting from the metaERG pipeline is available (45).

## Acknowledgments

Support for this research was provided in part by the National Institute of Health award (CA205932) to A.E.M., B.J.B., and P.S.G.C., with additional support from National Science Foundation awards (OPP-0442857, ANT-0838776, and PLR-1341339) to B.J.B., and Desert Research Institute (Institute Project Assignment) to A.E.M..

The assistance of several collaborators and students are acknowledged including C. Amsler, M. Amsler, L. Bishop, J. Cuce, B. Dent, N. Ernster, C. Gleasner, A. Maschek, A. Shilling, L. Siao, S. Thomas, and the Palmer Station science support staff. We especially would like to acknowledge the help of A. Oren in assisting with bacterial nomenclature proposed here. Likewise, we thank J.T. Hollibaugh and A.L. Reysenbach for comments on previous drafts of the manuscript.

## Supplemental Material

### Supplemental Materials and Methods

Details concerning ascidian sample collections, sample processing and high molecular weight DNA extraction, metagenome sequencing, metagenome binning, bin taxonomic and functional classification, Real Time PCR, manual assembly procedures, annotation and phylogenomic analyses and associated references are provided in the supplemental materials and methods text section.

### Supplemental Figure Legends

**Figure S1**. Ribosomal RNA-based phylogenetic tree. Sequences were selected from Verrucomicrobia (Phylum) and mostly Opitutaceae (family) genomes represented by isolates, metagenome assembled genomes and single cell assembled genomes from host associated and marine ecosystems. RAXML tree is based on 1,636 bases; 250 bootstraps were run, and values > 49 are shown at nodes. *Ca*. S. palmerolidicus (*Opitutaceae* bin 8) is indicated with an orange diamond symbol. Symbols designate environmental origins of the organisms: free-living are represented by circles: light blue - marine, green– freshwater, red – hydrothermal mud, brown – soils. Host-associated taxa from marine systems with blue diamonds, and from terrestrial systems with black diamonds.

**Figure S2**. Phylogenetic relationships of homologs of (a) MacB CDS and (b) celluase glycosylhydrolase family 5 enzymes identified in the Ca. S. palmerolidicus genome and closest neighboring sequences that were sourced from GenBank using BLAST. Maximum likelihood analysis was conducted using RAxML v. 8.2.12 using the PROTGAMMALG model with 550 bootstrap replicate trees calculated for (a) and 1000 for (b). Bootstrap values > 49 are shown which represent percent of times the topology was found.

**Figure S3**. Bacterial microcompartment (BMC) super loci identified in the *Ca*. S. palmerolidicus MAG. Conserved structural elements identified with Pfam and Swiss-Prot annotations were consistent with other PV-BMC loci which are associated carbohydrate utilization domains.

Transcriptional regulatory genes are indicated in green, conserved PV-BMC enzymes in blue, the BMC proteins in red, enzymes associated with rhamnose metabolism in violet and those with lactate utilization in purple.

### Supplemental Table Legends

**Table S1**. Metagenome sequencing metadata for *S. adareanum* microbiome. Host lobe identities indicate sample site (Nor, Norsel, Bon, Bonaparte Point, Del, DeLaca Island; followed by lobe designation, and year sampled). Assembly 1 joined two data sets using mega-merged contigs from 454 and Ion Proton sequence data sets. Assembly 2 included Assembly 1 and PacBio assemblies in addition to the manually assembled *Ca*. S. palmerolidicus MAG.

**Table S2**. Assembly 1 antiSMASH results for the 102 contigs > 40 Kbp. The 72.1 and 78.1 regions encoded biosynthetic gene clusters presented two putative clusters that were further investigated using QPCR to target samples with high BGC copy numbers for a subsequent round of metagenome sequencing.

**Table S3**. Quality checks and taxonomic classification using CheckM and GTDB-Tk for Opitutaceae-associated bins at different stages of analysis (most recent, highest quality on the left-most column and initial Coassembly 1 and bin 4 in the right-most column). The *Ca*. S. palmerolidicus-2 column reflects identifications (MetaERG annotation) of many of the missing markers in *Ca*. S palmerolidicus-1. The Opitutaceae bin 8 – clean CoAssembly 2 data set reflects an effort to remove non-Opitutaceae sequences from the bin (see supplemental material, methods). CoAssembly 1 bins 1 and 2 included the palmerolide A biosynthetic gene cluster and were otherwise populated by short contigs, and were taxonomically unidentified. CoAssembly 1 bin 4 included Opitutales-related contigs including a SSU rRNA gene sequence.

**Table S4**. GC percent for Opitutaceae-family representatives. Different genera are represented with different colors. Marine Optitutaceae metagenome assembled genomes are indicated with *. Source and species nomenclature: GTDB release 05-RS95. The GC content for *Ca*.

Synoicihabitans palmerolidicus is 58.7%.

**Table S5**. Quantitative PCR gene targets and primers. Primers targeting three gene targets were used in this study Primer IDs are based on bases from the beginning of the CDS. A GBlocks synthetic DNA positive control was used for estimating copy number^†^.

**Table S6**. Conserved markers used in phylogenomic analysis. Integrated microbial genomes (IMG) and/or GenBank accession numbers are listed. Shaded markers were those used for phylogenomic analysis (Fig. 4).

