## Supplementary material for "Discovery of an Antarctic ascidian-associated uncultivated *Verrucomicrobia* with antimelanoma palmerolide biosynthetic potential": All Supplemental Material

#### Materials and Methods

**Sample collections.** *Synoicum adareanum* lobe samples selected for metagenome sequencing were collected by SCUBA and stored frozen in a buffer (50 mM Tris-HCl at pH 8.0, 50 mM EDTA pH 8.0, 15% sucrose; (1)) at -80 °C until DNA extraction. Samples (Nor-2a-2007 and Nor-2C-2007) were collected from Norsel Point (S 64° 45.638', W 64° 05.874') on 3 May 2007 at a depth of 29 m. Samples Bon1c-2011 and Del2b-2011 were also collected by SCUBA (2) from Bonaparte Point (24 Mar 2011 at 26.2 m) and Delaca Island (28 Mar 2011 at 22 m) and stored frozen (-80 °C) until DNA extraction (see (3) for further sample details).

**Sample processing and high molecular weight DNA extraction.** For Nor2a-2007 and Nor2C-2007 the outer tissue layer was removed using a scalpel, then ~2.5-gram tissue sections were manually homogenized (1 min.) using a sterile mortar (ice-cold) with a pestle in sterile seawater (12 mL). The cell suspension was sieved through 63 mm sterile Nitex mesh to remove large debris then transferred to a sterile 25 mL Oakridge tube (Nalgene) for centrifugation (300 x g, 15 min at 4 °C) to pellet large cells and debris. The supernatant was then centrifuged at 8000 x g to pellet the bacterial cells at 4 °C. The pellet was resuspended in 200 mL of buffer (50 mM Tris-HCl at pH 8.0, 50 mM EDTA (pH 8.0, 15% sucrose; (1)).

DNA was extracted using the Qiagen Blood and Tissue Kit (Qiagen, Inc.) following manufacturer's instructions. The gDNA was screened by PCR to verify low levels of host (eukaryote) contamination using primers 960F (GGC TTA ATT TGA CTC AAC RCG) and 1200R (5' GGG CAT CAC AGA CCT G 3';(4)). Ketosynthase gene amplification was confirmed in these gDNA extracts following Riesenfeld et al. (5).

Samples Bon-1c-2011 and Del-2b-2011 were collected and homogenized as described by (3), except larger sample sizes were used such that 2.35 and 2.15 grams of tissue (respectively) was homogenized using a MiniLys (Bertin-Instruments,Montigny-le-Bretonneux, France) in a 7 mL tube with 3 mL sterile NaCl (3.5%). High molecular weight gDNA extracted from cell preparations followed Massana et al., (1). Care was taken to preserve the high molecular weight nature of the material e.g., large bore pipet tips, Pasteur pipets used for organic extractions and HMW gDNA pellet was rehydrated at 2 °C overnight gently shaking following ethanol precipitation. These extracts were reprecipitated in 3M NaCl and ethanol (200 proof). Extracts were checked on 0.7 % agarose gels and quantified using a ND-1000 Nanodrop spectrophotometer (Thermo Fisher Scientific, Waltham MA). Subsamples from these same lobes were used for amplicon sequencing (Illumina Inc., San Diego, CA) of the variable 4-5 region of the rRNA gene (3).

**454 and Proton metagenome sequencing.** 454 pyrosequencing (performed at the Roy J Carver Biotechnology Center, University of Illinois at Urbana-Champaign) was conducted with a bacterial enriched metagenomic DNA preparation from *S. adareanum* lobe (Nor-2c-2007). The MiniLys metagenomic DNA was used to generate single-stranded DNA libraries and emulsion PCR according to established protocols (454 Life Sciences, Roche). Amplified library fragments were sequenced on a Roche Genome Sequencer FLX system initially with a titration sequencing run, followed by a full plate run. Next, an Ion Proton System (Ion Torrent; run at the Nevada

Genomics Center following manufacturer's library preparation and sequencing protocols) was used to sequence a metagenomic DNA sample prepared from *S. adareanum* lobe Nor-2a-2007. See materials and methods in the main manuscript for CoAssembly1 details.

**Pacific Biosciences sample preparation and metagenome sequencing details.** DNA quality control for samples Bon1c-2011 and Del2b-2011 was confirmed with Qubit Fluorometer (Invitrogen, Inc., Carlsbad CA), NanoDrop 1000 and pulsed field gel electrophoresis (PFGE; BioRad, Hercules, CA). To note, although high molecular weight quality was confirmed on the PFGE, the NanoDrop ratios of 260:230 ratios were 1.2-1.4, much lower than 1.8 which is recommended by Pacific Biosciences (PB; San Diego, CA). The 260/280 ratio improved significantly after two rounds of purification with AMPure® PB beads (initial step in the PB SMRTbell protocol), the 260/230 ratio remained low. gDNA was sheared in G-Tubes (Covaris, Woburn, MA) and purified with 0.45x volume of AMPure PB beads. SMRTbell (PacBio) libraries were prepared according to the PacBio protocol specified in "Procedure and Checklist-Preparing gDNA Libraries Using the SMRTbell Express Template Preparation Kit 2.0". The removal of single-strand overhangs was followed by DNA damage repair reaction, end repair/A-tailing reaction and overhang SMRTbell adapter ligation, with all the steps performed subsequently in one tube. After 0.45x volume AMPure PB purification Bon-1c-2011 and Del-2b-2011 SMRTbell libraries were size selected on Blue Pippin instrument with 6 Kbp and 5 Kbp lower cutoff respectively. Sequencing primer 4 was annealed and DNA polymerase 3.0 was bound to the templates. The libraries were sequenced on a Sequel (PB) with sequencing chemistry 3.0 and 10 or 20hr movies A total of 2 SMRT cells were sequenced for Bon-1c-2011 with a total data output of 23.7 Gb. A total of 4 SMRT cells were sequenced for Del-2b-2011 with a total data output of 4.3 Gb. We obtained 48,298 and 9,576 CCS reads from Bon-1C-2011 and Del-2B-2011, respectively. The average read length was 11,870 bp for Bon-1C-2011 and 10,491 bp for Del-2B-2011 CCS reads.

**Binning and bin taxonomic and functional classification.** We initially used MaxBin (6) to bin CoAssembly 1 based on the coverage depth, tetranucleotide frequencies and single-copy marker genes, which resulted in 20 "genome"-like bins, containing 63,218 contigs (73.2% of the assembly). CheckM (7) v1.0.11 was used to estimate the genome completeness and potential contamination based on conserved marker genes and perform taxonomic evaluation of the CoAssembly 1 bins using similarity of genomic characteristics, and proximity within a reference genome tree. GTDB-Tk v0.1.3 was also used to evaluate the binned contigs with respect to taxonomic classifications, based on alignment of concatenated marker genes and maximum-likelihood placement within a reference tree, its relative evolutionary divergence, and ANI to reference genomes from GTDB taxonomy database (8).

Initial binning of CoAssembly 1 resulted in 3 bins of interest (Table S3 in which the putative BGC contigs (Fig. 1) that were found in two taxonomically unresolved bins (Bin 1 and Bin 2) dominated by short contigs encoding mostly hypothetical genes with no taxonomic affiliation. Then a third bin was identified (Bin 4, 27 contigs, 143 Kbp) with several contigs attributable to Opitutales, and many additional unclassified contigs. Despite additional re-assemblies, we were not able to link the BGC with these Opitutales scaffolds, thus motivating another round of metagenome sequencing using Pacific Biosciences Sequel Systems technology (PacBio).

CoAssembly 2 contigs were binned using MaxBin2 (9). The bin quality was assessed using CheckM v1.1.2, and GTDB-Tk v1.0.2, and then was used for taxonomic classification of the bins as above. We implemented a bin-cleaning strategy prior to functional classification of the Opitutaceae bin 8 sequence to reduce errors in classification and assessment of functional properties contributed by contaminating contigs which were evident upon visualization tools provided by MetaERG. This strategy takes a conservative approach (i.e., there is some chance that contigs that were true Verrucomicrobia contigs were discarded), however we felt this was the most robust approach. Examination of those ORFs in contigs discarded suggested 5 out of 24 were classified as having some percentage of ORF assigned to an Opitutaceae genus. First, we used the GTDB-assigned taxonomy as the basis for classification. Next, a custom script was developed to screen contigs in bin 8 with a majority rules algorithm to retain them in the bin if the majority of ORFs on a given contig were assigned to the classified taxonomy. All contigs with the verified identity were placed in the "cleaned" bin (Table S3). The cleaned bin was then run through CheckM v1.1.2 and GTDB-Tk v1.3.0 (10) to evaluate the efficacy of the cleaning algorithm. This effort reduced contamination in the bin, yet retained a similar level of markers identified for Verrucomicrobia. Following manual assembly of the Opitutaceae genome, a re-run with CheckM suggested a number of markers were still absent, these were identified however, through inspection of the MetaERG annotation. Thus, nearly all markers were identified, resulting in a CheckM completeness estimate of 96.04 %. This may have been the result of poor representation of Verrucomicrobia genomes in the CheckM database (12 genomes in the 2015 database that is currently being updated) when doing orthologous searches. When classified using GTDB-Tk the results suggested the closest affiliate was an *Opitutaceae* MAG UBA6669 (the only genome in this un-named genus) – however the result was based on a low average nucleotide identity (75.26) with this medium quality MAG and a low GTDB-Tk alignment fraction (AF) score of 0.02 (calculated as sum of lengths of bidirectional best hits divided by sum of lengths of all genes in each genome separately; AF being most valuable only when circumscribing species).

**Real Time PCR.** Primer 3 (11), plugin to Geneious (Auckland, NZ) was used to design primers to three coding regions along the candidate palmerolide A BGC (non-ribosomal peptide synthase, acyltransferase, and 3-hydroxymethylglutaryl coenzyme A synthase; amplicon size of 120 bases ea.) for real time PCR following design and optimization criteria recommended by (12). Homology of the primers to sequences other than their targets was evaluated by BLAST to the metagenome assembly. A single GBlocks synthetic positive control was designed with all three gene targets (Integrated DNA Technologies, IDT, Coralville, IA, USA). The GBlocks control also included a 120 base control region matching a putative luciferase CDS (in the putative BGC) that was not used for quantitative assays (Table S5).

As described in the main text, a *S. adareanum* DNA sample set (n=63 *S. adareanum* lobes from 21 colonies) with high levels of palmerolide A were screened with the real time PCR Real time PCR assays on a Quant Studio 3 (Thermo Fisher Scientific, Inc.) at the Nevada Genomics Center. Reactions (15 mL) were run with Power SYBR® Green PCR Master Mix (Applied Biosystems, Thermo Fisher Scientific), following the manufacturer's protocol and thermal cycling conditions (Initial hold at 95 C for 10 minutes followed by 40 cycles of denaturation at 95 °C for 15 sec, and anneal/extension at 60 °C for 1 min. This annealing temperature was confirmed to produce optimal results for the three gene targets at a primer concentration of 0.3 M. Results were analyzed using QuantStudio™ Design and Analysis Software v1.4.3 (Thermo Fisher Scientific) in which gene target copy numbers per ng of DNA template were estimated from standard curves of the synthetic positive control. All reactions had high efficiencies (ave.

99.54, 1.31 s.d., n=13) and $r^2$ (ave. 0.997, 0.004 s.d., n=13). Pearson correlation coefficients were determined (Microsoft® Excel for Mac v. 16.16.24), then for gene target levels compared to palmerolide A concentrations determined by LC-MS and 16S rRNA gene (variable region 3-4) amplicon sequence variant occurrence levels reported in Murray et al. (3), and the data was plotted using SigmaPlot (v. 14; Systat, San Jose, CA, USA).

**Manual MAG assembly and annotation**. A manual approach was implemented to arrive at assembly of the *Opitutaceae* MAG of interest. Four bins from different assemblies of the CCS reads, that had 58% GC content (targeted GC percentage of the palmerolide A BGC), were assembled with phrap (overlap based assembler; (13, 14)) and the assembly was visualized with Consed (15). The CCS reads were used to close gaps and verify repetitive elements. A total of ten contigs were resolved, five of which corresponded to sequences that were similar to one another suggesting they were a form of repeated elements within the genome. Rigorous assessment of these repetitive elements, including linking each contig end to other contigs with read-pairing information, assessing estimated gap lengths, and reviewing read coverage along the contigs, strongly suggest that the ten contigs represent the complete genome where each of the five largely unique contig was flanked by contigs corresponding to the repeated elements.

The nature of the five repeated elements that encompass the palmerolide BGC (outlined in Fig. 1) is fully supported by read depth of coverage analysis (e.g., the portions of the palmerolide BGC that are inferred to be present in five copies have 5X the fold coverage of the unique sections of the genome). Due to the very large length of the repeated elements (36.1-73.9 Kbp), no long reads were identified that spanned the entire length of the repeated regions or sufficient amounts that would allow us to specifically order the ten contigs into a single scaffold. A visual representation of the genome in circular format was prepared in GCView (16) in which the five unique contigs, and one possible ordering of the palmerolide BGC repeats is displayed. We used MetaERG (17) and NCBI's PGAP pipeline upon MAG submission (18) as annotation pipelines for analysis of the palmerolide A-containing MAG.

**Phylogenomic analyses.** We targeted genomes associated with marine and host-associated habitats in the *Opitutaceae* family in addition to including representatives of all *Opitutaceae* genera represented in the GTDB (release 05-RS95). The 115 reference datasets were downloaded from the NCBI and JGI IMG databases. The genome sequences were annotated by Prokka v1.14.5 (19) which performed the open reading frame (ORF) calls and scanned the protein and domain databases in a hierarchical manner from the translated peptide sequences (see Table S6 for list of shared ribosomal proteins and rRNA genes).

Among 115 reference datasets, 62 (many assembled metagenomes and single cell genomes) included 16S rRNA sequences that were identified in the Prokka annotation result. Of these, 47 were unique 16S rRNA sequences without duplication (same genome multiple copy, or identical sequences). The *Opitutaceae* bin 8 (and assembled *Ca*. S. palmerolidicus genome) also included a 16S rRNA identified from the assembled genome. We added the previously sequenced 16S rRNA (FJ169192) and performed the multiple sequence by MUSCLE v3.8.31 (20) and resulting in 49 16S rRNA sequences with 1,636 aligned positions. A maximum likelihood tree was constructed using RAxML v.8.2.12 under the GTRCAT model of evolution and with the number of bootstraps automatically determined (MRE-based bootstopping criterion). A total of 250 bootstrap replicates were conducted under the rapid bootstrapping algorithm, with 100 sampled to generate proportional support values. The final tree is rooted by *Kiritimatiella* glycovorans L21-Fru-AB and displayed in MegaX (21).

There were 16 ribosomal proteins (*rplB*, *C, D, E, F, N, O, P, and rpsC, H, J, K, L, Q, M, S*) identified shared among 48 reference datasets. Each individual gene set was aligned using MUSCLE. The 16 alignments were concatenated, forming a final alignment comprising 48 genomes and 3,035 amino-acid positions. A maximum likelihood tree was constructed using RAxML v.8.2.12 (22) under the LG plus gamma model of evolution (PROTGAMMALG in the RAxML model section), and with the number of bootstraps automatically determined (MRE-based bootstrapping criterion). A total of 150 bootstrap replicates were conducted under the rapid bootstrapping algorithm, with 100 sampled to generate proportional support values. The final tree is rooted by *Kiritimatiella* glycovorans L21-Fru-AB, a distinct phylum-level lineage originally designated *Verrucomicrobia* (23) and displayed in MegaX.

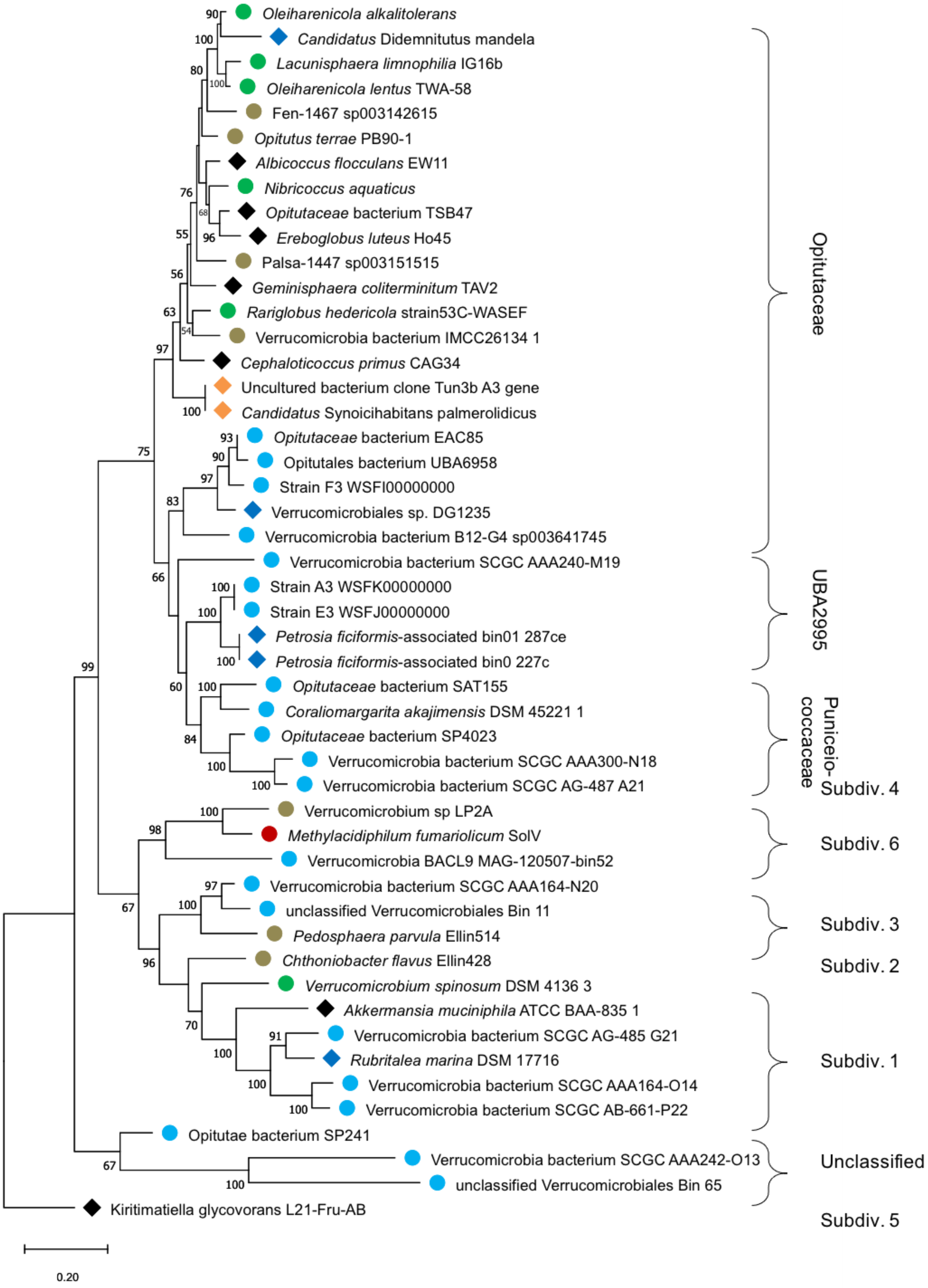
Oleiharenicola alkalitolerans
Candidatus Didemnitutus mandela
Lacunisphaera limnophilia IG16b
Oleiharenicola lentus TWA-58
Fen-1467 sp003142615
Opitutus terrae PB90-1
Albicoccus flocculans EW11
Nibricoccus aquaticus
Opitutaceae bacterium TSB47
Ereboglobus luteus Ho45
Palsa-1447 sp003151515
Geminisphaera coliterminitum TAV2
Rariglobus hedericola strain53C-WASEF
Verrucomicrobia bacterium IMCC26134 1
Cephaloticoccus primus CAG34
Uncultured bacterium clone Tun3b A3 gene
Candidatus Synoicihabitans palmerolidicus
Opitutaceae bacterium EAC85
Opitutales bacterium UBA6958
Strain F3 WSFI00000000
Verrucomicrobiales sp. DG1235
Verrucomicrobia bacterium B12-G4 sp003641745
Verrucomicrobia bacterium SCGC AAA240-M19
Strain A3 WSFK00000000
Strain E3 WSFJ00000000
Petrosia ficiformis-associated bin01 287ce
Petrosia ficiformis-associated bin0 227c
Opitutaceae bacterium SAT155
Coraliomargarita akajimensis DSM 45221 1
Opitutaceae bacterium SP4023
Verrucomicrobia bacterium SCGC AAA300-N18
Verrucomicrobia bacterium SCGC AG-487 A21
Verrucomicrobium sp LP2A
Methylacidiphilum fumariolicum SolV
Verrucomicrobia BACL9 MAG-120507-bin52
Verrucomicrobia bacterium SCGC AAA164-N20
unclassified Verrucomicrobiales Bin 11
Pedosphaera parvula Ellin514
Chthoniobacter flavus Ellin428
Verrucomicrobium spinosum DSM 4136 3
Akkermansia muciniphila ATCC BAA-835 1
Verrucomicrobia bacterium SCGC AG-485 G21
Rubritalea marina DSM 17716
Verrucomicrobia bacterium SCGC AAA164-O14
Verrucomicrobia bacterium SCGC AB-661-P22
Opitutae bacterium SP241
Verrucomicrobia bacterium SCGC AAA242-O13
unclassified Verrucomicrobiales Bin 65
Kiritimatiella glycovorans L21-Fru-AB
Opitutaceae
UBA2995
Puniceio-coccaceae
Subdiv. 4
Subdiv. 6
Subdiv. 3
Subdiv. 2
Subdiv. 1
Unclassified
Subdiv. 5
0.20

A

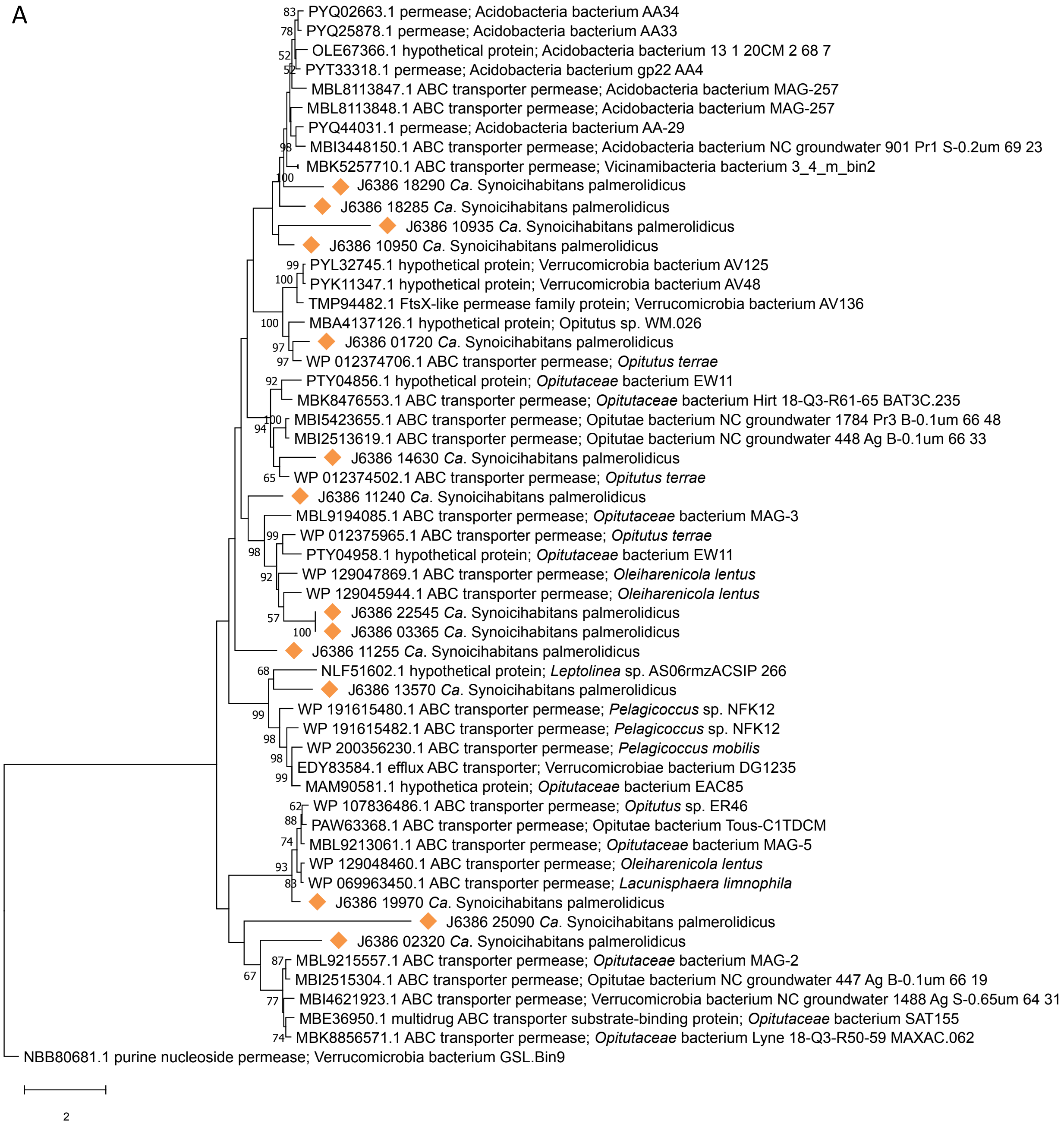

PYQ02663.1 permease; Acidobacteria bacterium AA34
PYQ25878.1 permease; Acidobacteria bacterium AA33
OLE67366.1 hypothetical protein; Acidobacteria bacterium 13 1 20CM 2 68 7
PYT33318.1 permease; Acidobacteria bacterium gp22 AA4
MBL8113847.1 ABC transporter permease; Acidobacteria bacterium MAG-257
MBL8113848.1 ABC transporter permease; Acidobacteria bacterium MAG-257
PYQ44031.1 permease; Acidobacteria bacterium AA-29
MBI3448150.1 ABC transporter permease; Acidobacteria bacterium NC groundwater 901 Pr1 S-0.2um 69 23
MBK5257710.1 ABC transporter permease; Vicinamibacteria bacterium 3_4_m_bin2
J6386 18290 *Ca*. Synoicihabitans palmerolidicus
J6386 18285 *Ca*. Synoicihabitans palmerolidicus
J6386 10935 *Ca*. Synoicihabitans palmerolidicus
J6386 10950 *Ca*. Synoicihabitans palmerolidicus
PYL32745.1 hypothetical protein; Verrucomicrobia bacterium AV125
PYK11347.1 hypothetical protein; Verrucomicrobia bacterium AV48
TMP94482.1 FtsX-like permease family protein; Verrucomicrobia bacterium AV136
MBA4137126.1 hypothetical protein; Opitutus sp. WM.026
J6386 01720 *Ca*. Synoicihabitans palmerolidicus
WP 012374706.1 ABC transporter permease; *Opitutus terrae*
PTY04856.1 hypothetical protein; *Opitutaceae* bacterium EW11
MBK8476553.1 ABC transporter permease; *Opitutaceae* bacterium Hirt 18-Q3-R61-65 BAT3C.235
MBI5423655.1 ABC transporter permease; Opitutae bacterium NC groundwater 1784 Pr3 B-0.1um 66 48
MBI2513619.1 ABC transporter permease; Opitutae bacterium NC groundwater 448 Ag B-0.1um 66 33
J6386 14630 *Ca*. Synoicihabitans palmerolidicus
WP 012374502.1 ABC transporter permease; *Opitutus terrae*
J6386 11240 *Ca*. Synoicihabitans palmerolidicus
MBL9194085.1 ABC transporter permease; *Opitutaceae* bacterium MAG-3
WP 012375965.1 ABC transporter permease; *Opitutus terrae*
PTY04958.1 hypothetical protein; *Opitutaceae* bacterium EW11
WP 129047869.1 ABC transporter permease; *Oleiharenicola lentus*
WP 129045944.1 ABC transporter permease; *Oleiharenicola lentus*
J6386 22545 *Ca*. Synoicihabitans palmerolidicus
J6386 03365 *Ca*. Synoicihabitans palmerolidicus
J6386 11255 *Ca*. Synoicihabitans palmerolidicus
NLF51602.1 hypothetical protein; *Leptolinea* sp. AS06rmzACSIP 266
J6386 13570 *Ca*. Synoicihabitans palmerolidicus
WP 191615480.1 ABC transporter permease; *Pelagicoccus* sp. NFK12
WP 191615482.1 ABC transporter permease; *Pelagicoccus* sp. NFK12
WP 200356230.1 ABC transporter permease; *Pelagicoccus mobilis*
EDY83584.1 efflux ABC transporter; Verrucomicrobiae bacterium DG1235
MAM90581.1 hypothetica protein; *Opitutaceae* bacterium EAC85
WP 107836486.1 ABC transporter permease; *Opitutus* sp. ER46
PAW63368.1 ABC transporter permease; Opitutae bacterium Tous-C1TDCM
MBL9213061.1 ABC transporter permease; *Opitutaceae* bacterium MAG-5
WP 129048460.1 ABC transporter permease; *Oleiharenicola lentus*
WP 069963450.1 ABC transporter permease; *Lacunisphaera limnophila*
J6386 19970 *Ca*. Synoicihabitans palmerolidicus
J6386 25090 *Ca*. Synoicihabitans palmerolidicus
J6386 02320 *Ca*. Synoicihabitans palmerolidicus
MBL9215557.1 ABC transporter permease; *Opitutaceae* bacterium MAG-2
MBI2515304.1 ABC transporter permease; Opitutae bacterium NC groundwater 447 Ag B-0.1um 66 19
MBI4621923.1 ABC transporter permease; Verrucomicrobia bacterium NC groundwater 1488 Ag S-0.65um 64 31
MBE36950.1 multidrug ABC transporter substrate-binding protein; *Opitutaceae* bacterium SAT155
MBK8856571.1 ABC transporter permease; *Opitutaceae* bacterium Lyne 18-Q3-R50-59 MAXAC.062
NBB80681.1 purine nucleoside permease; Verrucomicrobia bacterium GSL.Bin9
2

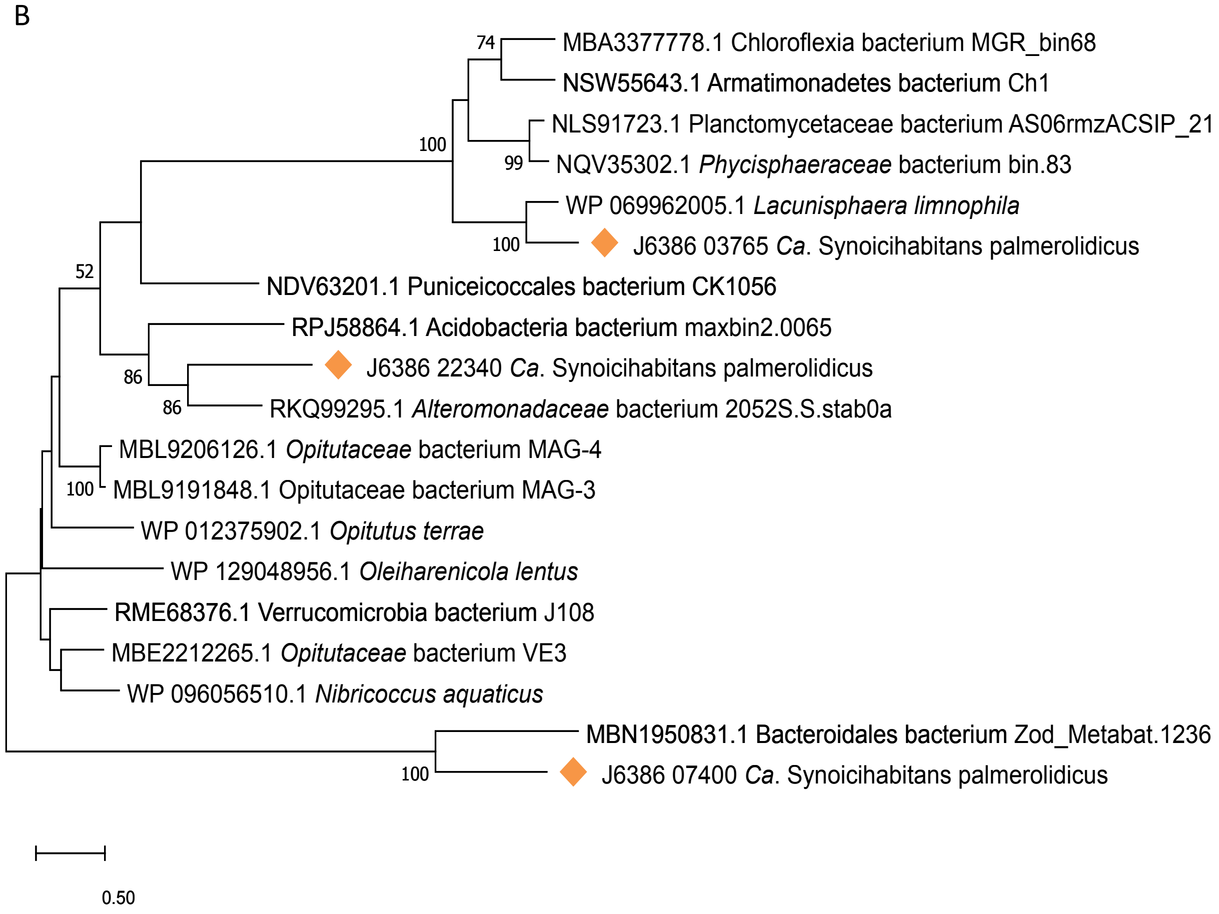
B
74
MBA3377778.1 Chloroflexia bacterium MGR_bin68
NSW55643.1 Armatimonadetes bacterium Ch1
NLS91723.1 Planctomycetaceae bacterium AS06rmzACSIP_21
100
99
NQV35302.1 Phycisphaeraceae bacterium bin.83
WP 069962005.1 Lacunisphaera limnophila
100
J6386 03765 Ca. Synoicihabitans palmerolidicus
52
NDV63201.1 Puniceicoccales bacterium CK1056
RPJ58864.1 Acidobacteria bacterium maxbin2.0065
86
J6386 22340 Ca. Synoicihabitans palmerolidicus
86
RKQ99295.1 Alteromonadaceae bacterium 2052S.S.stab0a
MBL9206126.1 Opitutaceae bacterium MAG-4
100
MBL9191848.1 Opitutaceae bacterium MAG-3
WP 012375902.1 Opitutus terrae
WP 129048956.1 Oleiharenicola lentus
RME68376.1 Verrucomicrobia bacterium J108
MBE2212265.1 Opitutaceae bacterium VE3
WP 096056510.1 Nibricoccus aquaticus
MBN1950831.1 Bacteroidales bacterium Zod_Metabat.1236
100
J6386 07400 Ca. Synoicihabitans palmerolidicus
0.50

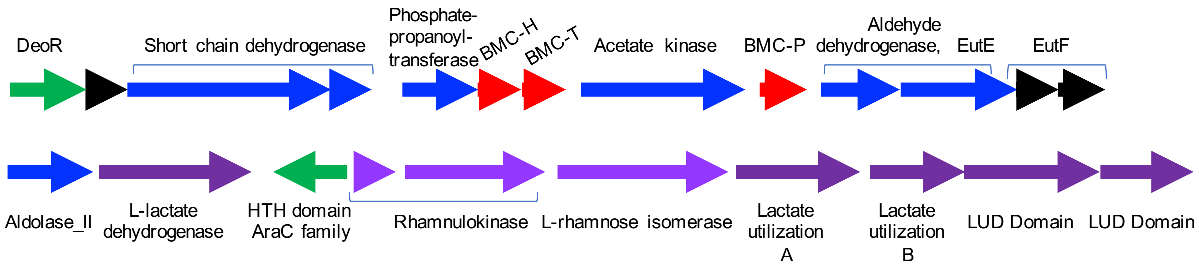
DeoR
Short chain dehydrogenase
Phosphate-
propanoyl-
transferase
BMC-H
BMC-T
Acetate kinase
BMC-P
Aldehyde
dehydrogenase, EutE
EutF
Aldolase_II
L-lactate
dehydrogenase
HTH domain
AraC family
Rhamnulokinase
L-rhamnose isomerase
Lactate
utilization
A
Lactate
utilization
B
LUD Domain
LUD Domain

| Metagenome samples | *Synoicum adareanum* lobe ID | | | | Metagenome Assemblies | |
|---|---|---|---|---|---|---|
| | Nor2c-2007 | Nor2a-2007 | Bon1c-2011 | Del2b-2011 | Nor2c-2007 and Nor2a2007 | Nor2c-2007, Nor2a2007, Bon1c-2011, Del2b-2011 |
| Sequencing Technology | 454 | Ion Proton | PacBio x 2 lanes | PacBio x 4 lanes | 454+Ion Proton | 454, Ion Proton, PacBio (Feb 2020) |
| Raw Reads | | | | | | |
| Reads | 1,570,137 | 89,330,870 | 2,499,689 | 3,014,737 | 90,900,996 | 5,600,821 |
| Total Bases (bp) | 904,455,285 | 17,053,251,055 | 23,936,900,178 | 4,127,276,138 | 17,957,706,340 | 28,213,247,451 |
| Mean Read Length (bp) | 550 | 190 | 9575.95 | 1369.03 | 197.55 | 5037.34 |
| Assembly | | | | | CoAssembly 1 | CoAssembly 2 |
| Number of contigs | - | - | 3,485 | 1,108 | 86,387 | 4,215 |
| N50 bp | - | - | 42,473 | 31,254 | 2,778 | 40,098 |
| Max contig size bp | - | - | 1,040,994 | 1,419,724 | 153,680 | 2,235,039 |
| Min contig size bp | - | - | 1,967 | 426 | 200 | 239 |
| total assembly size bp | - | - | 88,595,314 | 20,158,880 | 144,953,904 | 97,970,181 |

| Region | Type | From | To | Most similar cluster | Class | Similarity |
|---|---|---|---|---|---|---|
| 12.1 | Bacteriocin | 11,851 | 22,753 | | | |
| 16.1 | NRPS | 1 | 49,947 | nematophin | NRP | 25% |
| 92.1 | NRPS | 1 | 48,177 | Puwainaphycin F / minutissamide B, C, D | NRP | 61% |
| 24.1 | NRPS, T1PKS | 1 | 41,119 | rakicidin A / rakicidin B | NRP:Cyclic depsipeptide+polyketide: modular type 1 | 22% |
| 27.1 | NRPS, T1PKS | | 41,399 | | | |
| 38.1 | NRPS, transAT-PKS-like | 1 | 41,662 | | | |
| 84.1 | Terpene | 24,732 | 46,436 | hopene | Terpene | 23% |
| 87.1 | Terpene | 20,623 | 41,480 | | | |
| 72.1 | transAT-PKS-like, NRPS | 1 | 42,310 | chlorotonil A | Polyketide | 15% |
| 78.1 | transAT-PKS-like, NRPS | 1 | 68,977 | pyxipyrrolone A/ pyxipyrrolone B | NRP + Polyketide | 33% |

| Source | Bin Id | *Ca.* S. palmerolidicus-2 | *Ca.* S. palmerolidicus-1 | Bin8 - clean CoAssmembly 2 | Bin 8 CoAssmembly 2 | Bin 4 CoAssembly 1 | Bin 1 Coassembly 1 | Bin 2 Coassembly 1 |
|---|---|---|---|---|---|---|---|---|
| GTDB-Tk | Domain | Bacteria | Bacteria | Bacteria | Bacteria | Bacteria | Bacteria | Root (UID1) |
| | Phylum | Verrucomicrobiota | Verrucomicrobiota | Verrucomicrobiota | Verrucomicrobiota | Verrucomicrobiota | NA | NA |
| | Classification | Verrucomicrobiae | Verrucomicrobiae | Verrucomicrobiae | Verrucomicrobiae | Verrucomicrobiae | NA | NA |
| | Order | Opitutales | Opitutales | Opitutales | Opitutales | Opitutales | NA | NA |
| | Family | Optitutaceae | Optitutaceae | Optitutaceae | Optitutaceae | Optitutaceae | NA | NA |
| | Genus | UBA6669 | UBA6669 | UBA6669 | UBA6669 | NA | NA | NA |
| CheckM | # genomes | 88 | 88 | 88 | 88 | 88 | 5449 | 5656 |
| | # markers | 227 | 227 | 230 | 230 | 230 | 104 | 56 |
| | # marker sets | 146 | 146 | 148 | 148 | 148 | 58 | 24 |
| | 0 | 7 | 51 | 35 | 34 | 197 | 103 | 56 |
| | 1 | 218 | 174 | 184 | 169 | 33 | 1 | 0 |
| | 2 | 1 | 1 | 10 | 25 | 0 | 0 | 0 |
| | 3 | 0 | 0 | 1 | 2 | 0 | 0 | 0 |
| | 4 | 0 | 0 | 0 | 0 | 0 | 0 | 0 |
| | 5 | 1 | 1 | 0 | 0 | 0 | 0 | 0 |
| | Completeness | 96.04 | 71.89 | 80.86 | 82.04 | 4.59 | 1.72 | 0 |
| | Contamination | 3.42 | 3.42 | 4.62 | 8.43 | 0 | 0 | 0 |
| | Strain heterogeneity | 100 | 100 | 76.92 | 35.48 | 0 | 0 | 0 |
| GTDB-Tk | closest_placement_reference | NA | GCA_002455175.1 | GCA_002321895.1 | NA | NA | NA | NA |
| | closest_placement_radius | NA | 95 | 95 | NA | NA | NA | NA |
| | closest_placement_taxonomy - species | NA | UBA6669 sp002455175 | UBA6669 sp002455175 | NA | NA | NA | NA |
| | closest_placement_ani | NA | 75.26 | 75.16 | NA | NA | NA | NA |
| | closest_placement_af | NA | 0.02 | 0.02 | NA | NA | NA | NA |
| | classification_method | NA | taxonomic classification defined by topology and ANI | taxonomic classification defined by topology and ANI | NA | NA | NA | NA |
| | aa_percent | NA | 67.3 | 56.53 | 61.11 | NA | NA | NA |
| | red_value | NA | 0.920899756 | 0.920064441 | 0.910022563 | NA | NA | NA |

| Species | Strain | Genome accession | % GC |
|---|---|---|---|
| *Cephaloticoccus capnophilus* | CV41 | RS_GCF_001580045.1 | 60.54% |
| *Cephaloticoccus primus* | CAG34 | RS_GCF_001580015.1 | 63.03% |
| *Cephalitcoccus* sp002367815* | | GB_GCA_002367815.1 | 54.25% |
| *Cephalitcoccus* sp002713695* | | GB_GCA_002713695.1 | 55.96% |
| *Cephalitcoccus* sp002713695* | | GB_GCA_002471335.1 | 54.13% |
| *Ca*. Didemnitutus mandela* | | GB_GCA_002591725.1 | 51.93% |
| *Ereboglobus luteus* | Ho45 | RS_GCF_003096195.18 | 59.67% |
| *Ereboglobus* sp001650175 | TSB47 | RS_GCF_001650175.1 | 62.88% |
| *Geminisphaera colitermitum* | TAV2 | RS_GCF_000171235.2 | 60.93% |
| *Geminisphaera colitermitum* | TAV3 | RS_GCF_001653815.2 | 60.89% |
| *Geminisphaera colitermitum* | TAV4 | RS_GCF_001653825.2 | 60.85% |
| *Geminisphaera* sp000242935 | TAV5 | RS_GCF_000242935.2 | 63.45% |
| *Geminisphaera* sp000242935 | TAV1 | RS_GCF_000243495.5 | 63.25% |
| *Lacunisphaera limnophila* | IG16b | RS_GCF_001746835.1 | 66.48% |
| *Lacunisphaera* sp001464505 | | GB_GCA_001464505.1 | 64.27% |
| *Lacunisphaera* sp002304425 | | GB_GCA_002304425.1 | 56.80% |
| *Lacunisphaera* sp004118375 | TWA-58 | GB_GCA_004118375.1 | 65.30% |
| *Lacunisphaera* sp005787865 | | GB_GCA_005787865.1 | 56.64% |
| *Lacunisphaera* sp900104025 | GAS368 | GB_GCA_900104025.1 | 65.67% |
| *Nibricoccus aquaticus* | HZ-65 | RS_GCF_002310495.1 | 65.25% |
| *Opitutus terrae* | PB90-1 | RS_GCF_000019965.1 | 65.34% |
| *Opitutus* sp002288455 | | GB_GCA_002288455.1 | 67.69% |
| *Opitutus* sp002304435 | | GB_GCA_002304435.1 | 69.98% |
| *Opitutus* sp003054705 | ER46 | GB_GCA_003054705.1 | 67.07% |
| *Opitutus* sp005786775 | | GB_GCA_005786775.1 | 70.04% |
| *Opitutus* sp005792355 | | GB_GCA_005792355.1 | 64.74% |
| *Opitutus* sp005792355 | | GB_GCA_005779485.1 | 64.83% |
| B12-G4 sp003641745 | | GB_GCA_003641745.1 | 58.28% |
| DG1235 sp000155695 | DG1235 | RS_GCF_000155695.1 | 54.26% |
| EW11 sp003054665 | EW11 | RS_GCF_003054665.1 | 63.63% |
| Fen-1467 sp003142615 | | GB_GCA_003142615.1 | 64.26% |
| Fen-1467 sp003142615 | | GB_GCA_003141355.1 | 63.94% |
| Fen-1467 sp003142615 | | GB_GCA_003142335.1 | 64.50% |
| Fen-1467 sp003142955 | | GB_GCA_003142955.1 | 62.41% |
| Fen-1467 sp003142955 | | GB_GCA_003157795.1 | 62.27% |
| Fen-1467 sp003142955 | | GB_GCA_003154875.1 | 62.37% |
| Fen-1467 sp003153035 | | GB_GCA_003153035.1 | 64.04% |
| Fen-1467 sp003153035 | | GB_GCA_003160855.1 | 64.04% |
| Fen-1467 sp003153035 | | GB_GCA_003157495.1 | 64.27% |
| Fen-1467 sp003157575 | | GB_GCA_003157575.1 | 63.34% |
| IMCC26134 sp002382525 | | GB_GCA_002382525.1 | 63.52% |
| IMCC26134 sp002382525 | | GB_GCA_002383805.1 | 63.56% |
| IMCC26134_A sp000972765 | IMCC26134 | RS_GCF_000972765.1 | 61.28% |
| J108 sp003694825 | | GB_GCA_003694825.1 | 65.52% |
| Opi-474 sp003402695 | | GB_GCA_003402695.1 | 65.60% |
| Palsa-1447 sp003140665 | | GB_GCA_003140665.1 | 66.23% |

| Species | Strain | Genome accession | % GC |
|---|---|---|---|
| Palsa-1447 sp003140665 | | GB_GCA_003133585.1 | 66.47% |
| Palsa-1447 sp003140665 | | GB_GCA_003167995.1 | 66.10% |
| Palsa-1447 sp003140665 | | GB_GCA_003167115.1 | 66.40% |
| Palsa-1447 sp003151495 | | GB_GCA_003151495.1 | 66.51% |
| Palsa-1447 sp003151515 | | GB_GCA_003151515.1 | 67.03% |
| Palsa-1447 sp003164715 | | GB_GCA_003164715.1 | 66.99% |
| Palsa-1447 sp003168315 | | GB_GCA_003168315.1 | 66.69% |
| Tous-C4FEB sp002304445 | | GB_GCA_002304445.1 | 62.76% |
| Tous-C4FEB sp002304445 | | GB_GCA_002288405.1 | 62.94% |
| Tous-C4FEB sp002304445 | | GB_GCA_005789435 | 62.94% |
| Tous-C4FEB sp002737275 | | GB_GCA_002737275.1 | 62.84% |
| Tous-C4FEB sp005799525 | | GB_GCA_005799525.1 | 63.21% |
| UBA2377 sp002344205 | | GB_GCA_002344205.1 | 65.29% |
| UBA2377 sp002344205 | | GB-GCA_002343065.1 | 65.24% |
| UBA5691 sp002420185* | | GB_GCA_002320185.1 | 48.68% |
| UBA5691 sp002420265* | | GB_GCA_002420265.1 | 49.76% |
| UBA5691 sp002420265* | | GB_GCA_002722675.1 | 49.75% |
| UBA5691 sp002424265* | | GB_GCA_002170515.2 | 49.90% |
| UBA5691 sp002474325* | | GB_GCA_002474325.1 | 50.38% |
| UBA5691 sp002474325* | | GB_GCA_002450395.1 | 50.42% |
| UBA4691 sp002694885* | | GB_GCA_002694885.1 | 51.27% |
| UBA4691 sp002694885* | | GB_GCA_003525105.1 | 51.45% |
| UBA6669 sp002455175 | | GB_GCA_002455175.1 | 57.04% |
| | | *average* | 61.58% |
| | | *stdev* | 0.06 |

| Gene Target | Primer ID | Sequence |
|---|---|---|
| NRPS Condensation domain (NRPS) | PalA-NRPS _ 531F | 5'-CGCCAAGTTGGATCGAGACT-3' |
| | PalA-NRPS _ 650R* | 5'-GAGCGATTGGTGATTCCGGA-3' |
| Acyl transferase (AT) # | PalA-AT-1_624F | 5'-CACCGCACTACCCCATGAAT-3' |
| | PalA-AT-1_743R* | 5'-CAGACGGAACCTGAGTTCGT-3' |
| 3'hydroxymethylglutaryl synthase (HCS) | PalA-HCS_989F | 5'-ACTGGGTATTCGGCGTGAAG-3' |
| | PalA-HCS_1108R* | 5'-GGTGCGTAACTACCATCGGG-3' |

† GBlocks synthetic positive qPCR control sequence in which the NRPS positions 531-650 in the CDS are highlighted in yellow, luciferase positions 589-708 are highlighted in gray (not used in this study), AT1 positions 624-743 are highlighted in light blue and HCS are highlighted in green.

5’ATGTACTTGGAATCCGACACTTTTTTCGCCAAGTTGGATCGAGACTACTGGTTGCAGCGTTTCCCC
AAGGGGTTCCAGCCCGTGTTTCCCGCGAATGGCGATATTAAGGACTCCGGCGGGGAGATGGAGCG
ATTGGTGATTCCGGATTTTTTTCGACCTTGGAAGAACTCGCTAAAAAAATTGCGCGTTACCGTGCTGC
ACGGGCTGAGGCGGGCCTGGATCCGGTGGGAGGGTGTGTGACTCTCATGCTGCACACGTTTGTGG
ATCCCGATTTTTTTCACCGCACTACCCCATGAATCGACGACGCTAAAACAGATGTTGGCGCGGCAAA
TTACGAGCCCGGTGCGATGGACAGAGACCATGCGCTGGCTGTTTCGGCAGACGGAACCTGAGTTCG
TTTTTTTACTGGGTATTCGGCGTGAAGGATCAGGAAGTGGACTTAGCTCCTTACCGAAACCTCTACG
ACCAAACCTTGGCGGGCCGTGGTTTGTTGGTGTTGAAAGGGGTGCGTAACTACCATCGGGAATATG
ACTGGAGTTGATAA 3’

* Reverse sequences are not reverse and complimented.

### The acyltransferase domain targeted was the first of two domains in the putative *pal* BGC as oriented with 5' being the NRP at the beginning of the cluster.

| IMG ID | GenBank Accession | Taxon ID | infB | lepA | pheS | rplB | rplC | rplD | rplE | rplF | rplK | rplN | rplO | rplP | rpsB | rpsC | rpsE | rpsG | rpsH | rpsI | rpsJ | rpsK | rpsL | rpsM | rpsQ | rpsS | 16S rRNA |
|---|---|---|---|---|---|---|---|---|---|---|---|---|---|---|---|---|---|---|---|---|---|---|---|---|---|---|---|
| | | *Candidatus* Synoicohabitans palmerolidicus | + | + | + | + | + | + | + | + | + | + | + | + | + | + | + | + | + | + | + | + | + | + | + | + | + |
| 647533243 | GCA_000155695 | *Verrucomicrobiales* sp. DG1235 | + | + | + | + | + | + | + | + | + | + | + | + | + | + | + | + | + | + | + | + | + | + | + | + | + |
| 2228664034 | GCA_000383755 | Verrucomicrobia bacterium SCGC AAA300-N18 (unscreened) | + | + | - | + | + | + | + | - | - | + | - | + | - | + | - | + | + | - | + | - | + | - | + | + | + |
| 2228664038 | GCA_000385315 | Verrucomicrobia bacterium SCGC AAA164-P11 (unscreened) | - | - | - | - | - | - | - | - | - | - | - | - | - | - | - | - | - | - | - | - | - | - | - | - | + |
| 2228664042 | GCA_000385235 | Verrucomicrobia bacterium SCGC AAA164-A08 (unscreened) | - | - | - | - | - | - | - | - | - | - | - | - | - | - | - | - | - | - | - | - | - | - | - | - | - |
| 2228664043 | GCA_000385255 | Verrucomicrobia bacterium SCGC AAA164-B23 (unscreened) | - | - | - | - | - | - | - | - | - | - | - | - | - | - | - | - | - | - | - | - | - | - | - | - | - |
| 2228664044 | GCA_000385275 | Verrucomicrobia bacterium SCGC AAA164-I21 (unscreened) | - | - | + | + | + | + | + | + | - | + | + | + | - | + | + | + | + | - | + | + | + | + | + | + | + |
| 2228664050 | GCA_000383735 | Verrucomicrobia bacterium SCGC AAA164-M04 (unscreened) | - | - | - | - | - | - | - | - | + | - | - | - | + | - | - | - | - | + | - | - | - | - | - | - | - |
| 2228664051 | GCA_000385295 | Verrucomicrobia bacterium SCGC AAA164-N20 (unscreened) | - | - | - | + | + | + | + | + | - | + | + | + | - | + | + | + | + | - | + | + | + | + | + | + | + |
| 2236347002 | GCA_000383715 | Verrucomicrobia bacterium SCGC AAA164-E04 (unscreened) | + | - | + | + | + | + | + | + | + | + | + | + | - | + | + | + | + | + | + | + | + | + | + | + | + |
| 2236347003 | GCA_000264625 | Verrucomicrobia bacterium SCGC AAA168-E21 | + | - | + | - | - | - | - | - | + | - | - | - | - | - | - | - | - | - | - | - | - | - | - | - | + |
| 2236347021 | GCA_000264645 | Verrucomicrobia bacterium SCGC AAA168-F10 | + | - | + | + | + | + | + | + | + | - | + | + | - | + | - | + | - | - | + | + | + | + | + | + | + |
| 2236661017 | GCA_000382665 | Verrucomicrobia bacterium SCGC AAA300-K03 (unscreened) | - | - | - | + | + | + | + | + | + | + | + | + | + | + | + | + | + | - | + | + | + | + | + | + | + |
| 2236661018 | GCA_000382685 | Verrucomicrobia bacterium SCGC AAA300-O17 (unscreened) | - | - | - | + | + | + | + | + | + | + | + | + | + | + | + | + | + | - | + | + | + | + | + | + | - |
| 2517572138 | GCA_000264585 | Verrucomicrobia bacterium SCGC AAA164-A21 (unscreened) | - | - | - | - | - | - | - | - | - | - | - | - | - | - | - | - | - | - | - | - | - | - | - | - | - |
| 2517572141 | GCA_000264625 | Verrucomicrobia bacterium SCGC AAA168-E21 (unscreened) | + | - | + | - | - | - | - | - | + | - | - | - | - | - | - | - | - | - | - | - | - | - | - | - | + |
| 2517572140 | GCA_000264605 | Verrucomicrobia bacterium SCGC AAA164-O14 (unscreened) | - | + | + | + | + | + | + | + | - | + | - | + | + | + | + | + | + | - | + | - | + | - | + | + | + |
| 2517572142 | | Verrucomicrobia bacterium SCGC AAA168-F10 (unscreened) | + | - | + | - | - | - | - | - | + | - | - | - | - | - | - | - | - | - | - | - | - | - | - | - | + |
| 2616645036 | | *D. pulchra* _bleached_metagenome_bin377 | + | + | + | + | + | + | + | + | + | + | + | + | + | + | + | + | + | + | + | + | + | + | + | + | - |
| 2634166657 | | Verrucomicrobia bacterium SCGC AC-661-N10 (unscreened) | - | + | - | - | - | - | - | - | - | - | - | - | - | - | - | - | - | - | - | - | - | - | - | - | + |
| 2634166686 | | Verrucomicrobia bacterium SCGC AD-105-D22 (unscreened) | - | - | + | + | - | - | + | + | - | + | + | + | - | + | + | - | + | - | - | + | - | + | + | + | + |
| 2634166709 | | Verrucomicrobia bacterium SCGC AD-265-E10 (unscreened) | - | - | - | - | - | - | - | - | - | - | - | - | - | - | - | - | - | - | - | - | - | - | - | - | - |
| 2634166715 | | Verrucomicrobia bacterium SCGC AAA240-M19 (unscreened) | + | + | + | + | - | + | + | + | - | + | + | + | + | + | + | - | + | + | - | + | - | + | + | + | + |
| 2634166718 | | Verrucomicrobia bacterium SCGC AAA242-O13 (unscreened) | - | - | - | - | - | - | - | - | - | - | - | - | - | - | - | - | - | - | - | - | - | - | - | - | + |
| 2634166725 | | Verrucomicrobia bacterium SCGC AB-606-A23 (unscreened) | - | - | - | + | + | + | + | + | - | + | + | + | - | + | - | + | + | - | + | + | + | + | + | + | - |
| 2634166742 | | Verrucomicrobia bacterium SCGC AB-661-L11 (unscreened) | - | - | - | - | - | - | - | - | - | - | - | - | - | - | - | - | - | - | - | - | - | - | - | - | - |
| 2634166743 | | Verrucomicrobia bacterium SCGC AB-661-P22 (unscreened) | - | - | - | - | - | - | - | - | + | - | - | - | + | - | - | - | - | - | - | - | - | - | - | - | + |
| 2634166791 | | Verrucomicrobia bacterium SCGC AC-312-P03 (unscreened) | - | - | - | - | - | - | - | - | - | - | - | - | - | - | - | - | - | - | - | - | - | - | - | - | - |
| 2651870083 | | unclassified Verrucomicrobiales Bin 11 | + | + | - | - | - | - | - | - | - | - | - | - | + | - | - | - | - | + | - | + | - | + | - | - | + |
| 2651870084 | | unclassified Verrucomicrobiales Bin 46 | - | - | - | - | - | - | - | - | + | - | - | - | + | - | - | - | - | + | - | + | - | + | - | - | - |
| 2651870085 | | unclassified Verrucomicrobiales Bin 30 | - | - | - | - | - | - | + | + | - | - | + | - | - | - | + | + | + | - | - | + | + | + | - | - | - |
| 2651870086 | | unclassified Verrucomicrobiales Bin 34 | + | - | - | - | - | - | + | + | - | + | + | + | + | + | + | + | + | - | - | + | + | + | + | - | - |
| 2651870087 | | unclassified Verrucomicrobiales Bin 65 | - | - | - | + | - | - | - | - | - | + | - | + | - | + | - | - | - | - | - | - | - | - | + | + | + |
| 2657245246 | | Verrucomicrobia bacterium SCGC AC-312_D05v2 (unscreened) | - | - | - | - | - | - | - | - | - | - | - | - | - | - | - | - | - | - | - | - | - | - | - | - | + |
| 2657245247 | | Verrucomicrobia bacterium SCGC AC-312_D05v2 (contamination screened) | - | - | - | - | - | - | - | - | - | - | - | - | - | - | - | - | - | - | - | - | - | - | - | - | + |
| 2657245291 | | Verrucomicrobia bacterium SCGC AC-337_K21 - v2 (unscreened) | - | - | - | - | - | - | - | - | - | - | - | - | - | - | - | - | - | - | - | - | - | - | - | - | + |
| 2657245292 | | Verrucomicrobia bacterium SCGC AC-337_K21 - v2 (contamination screened) | - | - | - | - | - | - | - | - | - | - | - | - | - | - | - | - | - | - | - | - | - | - | - | - | + |
| 2657245323 | | Verrucomicrobia bacterium SCGC AC-312_O22 - v2 (unscreened) | - | + | - | - | - | - | - | - | - | - | - | - | - | - | - | - | - | - | - | - | - | - | - | - | + |
| 2657245324 | | Verrucomicrobia bacterium SCGC AC-312_O22 - v2 (contamination screened) | - | - | - | - | - | - | - | - | - | - | - | - | - | - | - | - | - | - | - | - | - | - | - | - | - |
| 2663762693 | | Verrucomicrobia bacterium SCGC AC-312_N19v2 (unscreened) | - | + | - | - | - | - | - | - | - | - | - | - | - | - | - | - | - | - | - | - | - | - | - | - | + |
| 2684622978 | | Verrucomicrobia bacterium SCGC AG-542-N08 | - | - | + | - | - | - | - | - | - | - | - | - | + | - | - | - | - | - | - | - | - | - | - | - | + |
| 2684622991 | | Verrucomicrobia bacterium SCGC AG-542-I16 | + | - | - | - | - | - | - | - | - | - | - | - | - | - | - | - | - | - | - | - | - | - | - | - | - |
| 2693429924 | | Verrucomicrobia bacterium SCGC AC-337_A14 - v2 (unscreened) | - | - | - | - | - | - | - | - | - | - | - | - | - | - | - | - | - | - | - | - | - | - | - | - | + |
| 2693429925 | | Verrucomicrobia bacterium SCGC AC-337_A14 - v2 (contamination screened) | - | - | - | - | - | - | - | - | - | - | - | - | - | - | - | - | - | - | - | - | - | - | - | - | + |
| 2693429980 | | Verrucomicrobia bacterium SCGC AC-337_J20 - v2 (unscreened) | - | - | - | - | - | - | - | + | - | - | + | - | - | - | + | - | + | - | - | + | - | + | - | - | - |
| 2693429981 | | Verrucomicrobia bacterium SCGC AC-337_J20 - v2 (contamination screened) | - | - | - | - | - | - | - | - | - | - | - | - | - | - | - | - | - | - | - | + | - | - | - | - | - |
| 2695420979 | | unclassified Verrucomicrobiales bin 1 | - | - | + | + | + | + | + | + | - | - | + | - | + | - | + | - | + | + | + | + | - | - | - | + | - |
| 2695420980 | | unclassified Verrucomicrobiales bin 2 | - | - | - | + | - | - | - | - | - | - | - | - | + | - | - | - | - | - | - | - | - | - | - | + | - |

| IMG ID | GenBank Accession | Taxon ID | infB | lepA | pheS | rplB | rplC | rplD | rplE | rplF | rplK | rplN | rplO | rplP | rpsB | rpsC | rpsE | rpsG | rpsH | rpsI | rpsJ | rpsK | rpsL | rpsM | rpsQ | rpsS | 16S rRNA |
|---|---|---|---|---|---|---|---|---|---|---|---|---|---|---|---|---|---|---|---|---|---|---|---|---|---|---|---|
| 2700988675 | | Verrucomicrobia bacterium SCGC AC-337_A09v3 (contamination screened) | - | - | - | - | - | - | - | - | - | - | - | - | - | - | - | - | - | - | - | - | - | - | - | - | + |
| 2731639125 | | Verrucomicrobia bacterium SCGC AG-485_L14 (contamination screened) | - | - | + | - | - | - | - | - | - | - | - | - | - | - | - | - | - | - | - | - | - | - | - | - | - |
| 2731639127 | | Verrucomicrobia bacterium SCGC AG-485_G21 (contamination screened) | - | - | - | + | + | + | + | - | + | + | + | + | - | + | + | + | + | - | + | + | + | + | + | + | + |
| 2731639129 | | Verrucomicrobia bacterium SCGC AG-487_A21 (contamination screened) | - | - | + | - | - | - | - | - | - | - | - | - | + | - | - | - | - | - | - | - | - | - | - | - | + |
| 2731639131 | | Verrucomicrobia bacterium SCGC AG-487_G17 (contamination screened) | + | - | - | - | - | - | - | - | - | - | - | - | - | - | - | - | - | - | - | - | - | - | - | - | + |
| 2739367775 | | Verrucomicrobiales bacterium JGI 01_E13 (contamination screened) | - | + | - | + | + | + | - | - | + | - | - | - | - | - | - | + | - | - | + | - | + | - | - | + | - |
| 2786546549 | GCA_002726435 | Opitutae bacterium NP78 | - | - | + | + | + | + | + | + | - | + | + | + | - | + | + | + | + | + | + | + | + | + | + | + | - |
| 2786546550 | GCA_002716045 | Opitutae bacterium SP85 | + | + | + | + | + | + | + | + | + | + | + | + | - | + | + | + | + | + | + | + | + | + | + | + | - |
| 2786546551 | GCA_002722545 | Opitutae bacterium SP216 | + | + | + | + | + | + | + | + | + | + | + | + | + | + | + | + | + | + | + | + | + | + | + | + | - |
| 2786546552 | GCA_002721705 | Opitutae bacterium SP241 | + | + | + | + | + | + | + | + | + | + | + | + | + | + | + | + | + | + | + | + | + | + | + | + | + |
| 2786546553 | GCA_002713685 | Opitutae bacterium SAT156 | + | + | - | + | + | + | + | + | - | + | + | + | + | + | + | + | + | - | + | + | + | + | + | + | - |
| 2786546554 | GCA_002730975 | Opitutae bacterium NP117 | + | + | + | + | + | + | + | + | + | + | + | + | - | + | + | + | + | + | + | + | + | + | + | + | - |
| 2786546555 | GCA_002695645 | Opitutae bacterium EAC647 | + | + | + | + | + | + | + | + | + | + | + | + | - | + | + | + | + | - | + | + | + | + | + | + | - |
| 2786546556 | GCA_002686835 | Opitutae bacterium ARS76 | - | + | + | + | + | + | + | + | + | + | + | + | + | + | + | + | + | + | + | + | + | + | + | + | - |
| 2786546557 | GCA_002689665 | Opitutae bacterium ARS1007 | - | + | - | + | + | + | + | + | + | + | + | + | + | + | + | + | + | + | + | + | + | + | + | + | - |
| 2786546558 | GCA_002692605 | Opitutae bacterium IN42 | + | + | - | + | + | + | + | + | + | + | + | + | + | + | + | + | + | + | + | + | + | + | + | + | - |
| 2786546559 | GCA_002702485 | Opitutae bacterium NAT243 | - | - | - | + | + | + | + | + | + | + | + | - | - | + | + | + | + | + | + | + | + | + | + | + | - |
| 2786546560 | GCA_002694035 | Opitutae bacterium IN48 | + | + | + | + | + | + | + | + | + | + | + | + | + | + | + | + | + | + | + | + | + | + | + | + | - |
| 2786546561 | GCA_002692535 | Opitutae bacterium IN921 | + | + | - | - | - | - | + | + | + | + | + | + | + | + | + | - | + | + | - | + | - | + | + | + | - |
| 2786546708 | GCA_002725655 | Opitutales bacterium NP990 | - | + | + | + | + | + | + | + | + | + | + | + | - | + | + | + | + | - | + | + | + | + | + | + | - |
| 2786546709 | GCA_002724235 | Opitutales bacterium RS400 | + | + | + | - | - | - | - | - | - | - | - | - | - | - | - | - | - | + | - | - | - | - | - | - | - |
| 2786546710 | GCA_002720915 | Opitutales bacterium SP2995 | + | + | + | + | + | + | - | - | - | - | - | + | + | + | - | - | - | + | - | - | - | - | - | + | - |
| 2786546935 | GCA_002713695 | *Opitutaceae* bacterium SAT155 | + | + | + | + | + | + | + | + | + | + | + | + | + | + | + | + | + | + | + | + | + | + | + | + | + |
| 2786546936 | GCA_002694885 | *Opitutaceae* bacterium EAC85 | + | + | - | + | + | + | + | + | + | + | + | + | + | + | + | + | + | + | + | + | + | + | + | + | + |
| 2786546938 | GCA_002722675 | *Opitutaceae* bacterium SP211 | + | + | - | + | + | + | + | + | - | + | + | + | - | + | + | + | + | + | + | - | + | + | - | + | - |
| 2786546939 | GCA_002717045 | *Opitutaceae* bacterium SP4023 | + | + | + | + | + | + | + | + | + | + | + | + | + | + | + | + | + | + | + | - | + | + | + | + | + |
| | GCA_002591725 | *Candidatus* Didemnitutus mandela | + | + | + | + | + | + | + | + | + | + | + | + | + | + | + | + | + | - | + | + | + | + | + | + | + |
| | GCA_003525105 | Opituae bacterium UBA10075 | + | - | + | + | + | + | - | - | - | - | - | + | + | + | - | + | - | - | + | + | + | + | + | + | - |
| | GCA_003482665 | *Opitutaceae* bacterium UBA8745 | + | - | - | + | + | + | + | + | - | + | - | + | - | + | - | + | + | - | + | + | + | + | + | + | - |
| | GCA_002420265 | Opitutales bacterium UBA5691 | + | + | + | + | + | + | + | + | + | + | + | + | - | + | + | - | + | + | + | + | - | + | + | + | - |
| | GCA_002420185 | Opitutales bacterium UBA5694 | + | + | + | + | + | + | + | + | + | + | + | + | + | + | + | + | + | + | + | + | + | + | + | + | - |
| | GCA_002450395 | Opitutales bacterium UBA6958 | + | + | + | + | + | + | + | + | + | + | + | + | - | + | + | + | + | + | + | + | + | + | + | + | + |
| | GCA_002474325 | Opitutales bacterium UBA7389 | + | + | + | + | + | + | + | + | + | + | + | + | + | + | + | + | + | + | + | + | + | + | + | + | - |
| | GCA_002367815 | *Opitutaceae* bacterium UBA3033 | + | - | + | + | + | + | + | + | - | + | + | + | + | + | + | + | + | + | + | + | + | + | + | + | - |
| | GCA_002471335 | *Opitutaceae* bacterium UBA7327 | + | + | + | + | + | + | + | + | + | + | + | + | + | + | + | + | + | + | + | + | + | + | + | + | - |
| | GCA_000019965 | *Opitutus terrae* PB90-1 | + | + | + | + | + | + | + | + | + | + | + | + | + | + | + | + | + | + | + | + | + | + | + | + | + |
| | GCA_000025905 | *Coraliomargarita akajimensis* DSM 45221 | + | + | + | + | + | + | + | + | + | + | + | + | + | + | + | + | + | + | + | + | + | + | + | + | + |
| | GCA_000171235 | *Geminisphaera coliterminitum* TAV2 | + | + | + | + | + | + | + | + | + | + | + | + | + | + | + | + | + | + | + | + | + | + | + | + | + |
| | GCA_000172555 | *Pedosphaera parvula* Ellin514 | + | + | + | + | + | + | + | + | + | + | + | + | + | + | + | + | + | + | + | + | + | + | + | + | + |
| | GCA_000173075 | *Chthoniobacter flavus* Ellin428 | + | + | + | + | + | + | + | + | + | + | + | + | + | + | + | + | + | + | + | + | + | + | + | + | + |
| | GCA_000378105 | *Rubritalea marina* DSM 17716 | + | + | + | + | + | + | + | + | + | + | + | + | + | + | + | + | + | + | + | + | + | + | + | + | + |
| | GCA_000972765 | Verrucomicrobia bacterium IMCC26134 | + | + | + | + | + | + | + | + | + | + | + | + | + | + | + | + | + | + | + | + | + | + | + | + | + |
| | GCA_001017655 | *Kiritimatiella glycovorans* L21-Fru-AB | + | + | + | + | + | + | + | + | + | + | + | + | + | + | + | + | + | + | + | + | + | + | + | + | + |
| | GCA_001438005 | Verrucomicrobia BACL9 MAG-120507-bin52 | + | + | - | + | + | + | + | + | + | + | + | + | + | + | + | + | + | + | + | + | + | + | + | + | + |
| | GCA_001580015 | *Cephaloticoccus primus* CAG34 | + | + | + | + | + | + | + | + | + | + | + | + | + | + | + | + | + | + | + | + | + | + | + | + | + |
| | GCA_001650175 | *Opitutaceae* bacterium TSB47 | + | + | + | + | + | + | + | + | + | + | + | + | + | + | + | + | + | + | + | + | + | + | + | + | + |
| | GCA_001746835 | *Lacunisphaera limnophilia* IG16b | + | + | + | + | + | + | + | + | + | + | + | + | + | + | + | + | + | + | + | + | + | + | + | + | + |
| | GCA_000020225 | *Akkermansia muciniphila* ATCC BAA-835 | + | + | + | + | + | + | + | + | + | + | + | + | + | + | + | + | + | + | + | + | + | + | + | + | + |
| | GCA_000172155 | *Verrucomicrobium spinosum* DSM 4136 | + | + | + | + | + | + | + | + | + | + | + | + | + | + | + | + | + | + | + | + | + | + | + | + | + |
| | GCA_000526255 | *Verrucomicrobium* sp. LP2A | + | + | + | + | + | + | + | + | + | + | + | + | + | + | + | + | + | + | + | + | + | + | + | + | + |
| | GCA_000953475 | *Methylacidiphilum fumariolicum* SolV | + | + | + | + | + | + | + | + | + | + | + | + | + | + | + | + | + | + | + | + | + | + | + | + | + |
| 2786546940 | GCA_003054665 | *Albicoccus flocculans* EW11 | - | + | + | + | + | + | + | + | + | + | + | + | - | + | + | + | + | + | + | + | + | + | - | + | + |
| | GCA_002455175 | *Opitutaceae* bacterium UBA6669 | - | + | + | - | - | - | - | + | + | + | + | + | - | + | - | - | + | + | - | + | - | + | - | - | - |
| | GCA_002344205 | *Opitutaceae* bacterium UBA2377 | - | + | + | + | + | + | + | + | + | + | + | + | - | + | - | + | + | + | + | + | + | + | - | + | - |

| IMG ID | GenBank Accession | Taxon ID | infB | lepA | pheS | rplB | rplC | rplD | rplE | rplF | rplK | rplN | rplO | rplP | rpsB | rpsC | rpsE | rpsG | rpsH | rpsI | rpsJ | rpsK | rpsL | rpsM | rpsQ | rpsS | 16S rRNA |
|---|---|---|---|---|---|---|---|---|---|---|---|---|---|---|---|---|---|---|---|---|---|---|---|---|---|---|---|
| | GCA_003142615 | Fen-1467 sp003142615 | - | + | + | + | + | + | + | + | + | + | + | + | - | + | + | + | + | + | + | + | + | + | - | + | + |
| | GCA_002310495 | *Nibricoccus aquaticus* | - | + | + | + | + | + | + | + | + | + | + | + | - | + | + | + | + | + | + | + | + | + | - | + | + |
| | GCA_003096195 | *Ereboglobus luteus* Ho45 | - | + | + | + | + | - | + | + | + | + | + | + | - | + | - | + | + | + | + | + | + | + | - | + | + |
| | GCA_003641745 | B12-G4 sp003641745 | - | + | + | + | + | + | + | + | - | + | + | + | - | + | + | + | + | + | + | + | + | + | - | + | + |
| | GCA_002382525 | IMCC26134 sp002382525 | - | + | + | + | + | + | + | + | + | + | + | + | - | + | + | + | + | + | + | + | + | + | - | + | - |
| | GCA_003694825 | J108 sp003694825 | - | + | + | + | + | + | + | + | + | + | + | + | - | + | + | + | + | + | + | + | + | + | - | + | - |
| | GCA_003402695 | Opi-474 sp003402695 | + | + | + | + | + | + | + | + | + | + | + | + | - | + | - | + | + | + | + | + | + | + | - | + | - |
| | GCA_003151515 | Palsa-1447 sp003151515 | + | + | + | + | + | + | - | + | + | + | + | + | - | + | - | + | + | + | + | + | + | + | - | + | + |
| | GCA_002304445 | Tous-C4FEB sp002304445 | + | + | + | + | + | + | + | + | + | + | + | + | - | + | - | - | + | + | + | + | - | + | - | + | - |
| | JAABVE00000000( | *Petrosia ficiformis*- associated bin0 (227c) | - | + | + | + | + | + | + | + | + | + | + | + | - | + | + | + | + | - | + | + | + | + | + | + | + |
| | JAABVD00000000( | *Petrosia ficiformis*- associated bin01 (287ce) | + | + | + | + | + | + | + | + | + | + | + | + | - | + | + | + | + | - | + | + | + | + | + | + | + |
| | GCA_014529625.1 | Strain F3 | + | + | + | + | + | + | + | + | + | + | + | + | - | + | - | + | + | + | + | + | + | + | + | + | + |
| | GCA_014529675.1 | Strain E3 | + | + | + | + | + | + | + | + | + | + | + | + | + | + | + | + | + | + | + | + | + | + | + | + | + |
| | GCA_014529665.1 | Strain A3 | + | + | + | + | + | + | + | + | + | + | + | + | + | + | + | + | + | + | + | + | + | + | + | + | + |
